# A Computational Study of the Role of Counterions and Solvent Dielectric in Determining the Conductance of B-DNA

**DOI:** 10.1101/2023.03.29.534812

**Authors:** Yiren Wang, Busra Demir, Hashem Mohammad, Ersin Emre Oren, M.P. Anantram

## Abstract

DNA naturally exists in a solvent environment, comprised of water and salt molecules such as sodium, potassium, magnesium, etc. Along with the sequence, the solvent conditions become a vital factor determining DNA structure and thus its conductance. Over the last two decades, researchers have measured DNA conductivity both in hydrated and almost dry (dehydrated) conditions. However, due to experimental limitations (the precise control of the environment), it is very difficult to analyze the conductance results in terms of individual contributions of the environment. Therefore, modeling studies can help us to gain a valuable understanding of various factors playing a role in charge transport phenomena. DNA naturally has negative charges located at the phosphate groups in the backbone, which provides both the connections between the base pairs and the structural support for the double helix. Positively charged ions such as the Sodium-ion (Na^+^), one of the most commonly used counterions, balance the negative charges at the backbone. This modeling study investigates the role of counterions both with and without the solvent (water) environment on charge transport through double-stranded DNA. Our computational experiments show that in dry DNA, the presence of counterions affects electron transmission at the lowest unoccupied molecular orbital energies. However, in solution, the counterions have a negligible role in transmission. Using the polarizable continuum model calculations, we demonstrate that the transmission is significantly higher at both the highest occupied and lowest unoccupied molecular orbital energies in a water environment as opposed to in a dry one. Moreover, calculations also show that the energy levels of neighboring bases are more closely aligned to ease electron flow in the solution.

## I. INTRODUCTION

DNA is one of the leading materials in molecular electronics due to its self-assembly property and long-range charge transport [1]. It naturally exists in a solvent environment, surrounded by water and salt molecules. Depending on the sequence of DNA, the environmental factors, such as the dielectric constant of solvents, the position and/or the local density of counterions surrounding DNA, become essential in determining DNA conformation and thus conductance. The electronic properties of DNA have been actively studied over the last two decades [1– 18]. However, the complex environment that a DNA molecule is in makes it challenging to analyze and interpret experimental results. Thus, modeling methods of varying degrees of complexity and accuracy become vital to understanding features in DNA conductance.

In several experiments, DNA conductance has been measured in both hydrated and dehydrated conditions [9,19]. These studies cannot easily separate the solvent contribution to conductance as some solvent molecules remain attached to the DNA even in the dehydrated case [20–22]. Initially, theoretical studies of charge transport were performed in a dehydrated environment [23] due to high computational costs. More recently, studies have begun modeling the role of the solvent environment on charge transport [1,24,25]. Kubař et al. suggested that the onsite energies can change with time by about 0.4 eV due to fluctuations in the solvent [26]. The offdiagonal hopping parameters can, in turn, affect DNA structure and transmission. Further, some studies included the influence of conformation and the solvent to investigate their joint role in determining conductance [25,27]. Reference suggested that solvent effects can decrease the sequence dependence of conductance. On the other hand, recent papers [16,28] show that even a single base-pair mismatch can be detected experimentally in the presence of a solvent. Further, [29] analyzed the experimentally obtained values of conductance using a pure machine learning approach and concluded that the experimental data shows differences in the conductance of strands with a single mismatch.

Wolter et al. used ab initio molecular dynamics simulation and concluded that DNA in a micro-hydrated environment, which retains the structure of the close solvation shell, shows charge transport properties similar to fully solvated DNA. They disagreed with experimental studies suggesting a strong humidity dependence of DNA conductance by arguing that in the presence of high humidity/solvent content, the conductivity of the residual solvent (not DNA) is measured [24]. The early work of Berashevich et al. studied the influence of humidity on DNA by hydrating base pairs with water molecules attached to the DNA bases [30]. They found hydration increased the bandgap of bases and concluded that this would make the conductance sensitive to the water environment. The effect of solvent on the structural properties of DNA has also been studied computationally using molecular dynamics (MD) simulations [23,31–34]. These studies show that the interaction between DNA molecules and the solvent contributes to fluctuations, leading to changes in the energy of both molecular orbitals and ionization potentials. Previously, it also has been demonstrated that solvent environment affects DNA conformation [15,35–37]. The common B-form is found at neutral pH and normal saline [38], while the A-form prefers dry/dehydrated conditions. On the other hand, A-form has a shorter distance between the base pairs when compared to B-form. The base pairs of A-form DNA are located away from the helical axis and are closer to the major groove [38]. Based on the prior work summarized above, it is challenging to distinguish the role of solvent environment and conformation from one another in modeling studies. To only reveal the role of solvent, in this model study, we kept the DNA structure fixed and only changed the dielectric constant of the environment in our calculations.

Apart from the solvent, counterions are also present close to the sugar-phosphate backbone. Positively charged counterions (such as Na^+^) are attracted to the negative charge on the phosphate group of the backbone. Prior studies have considered the effect of the counterions and concluded that they also affect DNA conductance. However, it has been difficult to reach clear conclusions due to the plethora of techniques and approximations [1,24,27,39,40]. The rich diversity in conclusions reached depends on the experimental and modeling methods and calls for a continued systematic study of the problem. In this modeling study, we focus on investigating the roles of Na^+^ ions and the solvent environment in determining the intrinsic conductance through DNA. The importance of this model study is the results help understand the role of solvent and counterions alone in determining the transmission without convolving the structural effects. We use the textbook forms of B-DNA, a natural form in water, with the sequence of 5’-CCCGCGCCC-3’. We chose this sequence based on previously published work [15,41]. Our results show that Na^+^ ions can significantly impact the charge transport properties of the DNA strand depending on the dielectric constant of the environment. In the dehydrated condition (low dielectric constant), the addition of Na^+^ ions lowers the bandgap to 0.77 eV compared with water (high dielectric constant), which has a bandgap of 4.03 eV. This difference is because Na^+^ ions add unoccupied energy levels in the bandgap of the DNA in a dehydrated condition. In contrast, for the water solvent, Na^+^ counterions add unoccupied energy levels that have higher energy than the LUMO (lowest unoccupied molecular orbital), which is primarily located on the DNA. This observation can be attributed to the high dielectric constant of water, which reduces the interaction between DNA and Na^+^ ions due to the charge screening effect. To demonstrate the generalizability of our conclusions, we applied the same simulation procedure to various DNA sequences with different lengths. Please refer to Section III of Supplemental Material for detailed discussions.

In addition, we find that the high dielectric constant increases the electronic coupling between the molecular orbitals of the DNA and yields a smaller onsite energy separation between them. Therefore, the transmission is at least two orders of magnitude larger at HOMO (highest occupied molecular orbital) and LUMO regions of the DNA with the water solvent. The rest of this article is structured as follows. Section II discusses our simulation procedure, including DFT and charge transport calculations with Green’s function method. In section III we compare energy levels, transmission plots, wavefunction analysis, and hopping parameters of the DNA molecule in both water and dry cases. Finally, we summarize the concluding remarks in Section IV.

## II. METHODS

The simulation procedure can be broken down into three steps: obtaining the atomic coordinates of the DNA molecule and counterions, performing density functional theory (DFT) calculations, and calculating the transmission using Green’s function approach. (see FIG. 1).

**FIG. 1.**
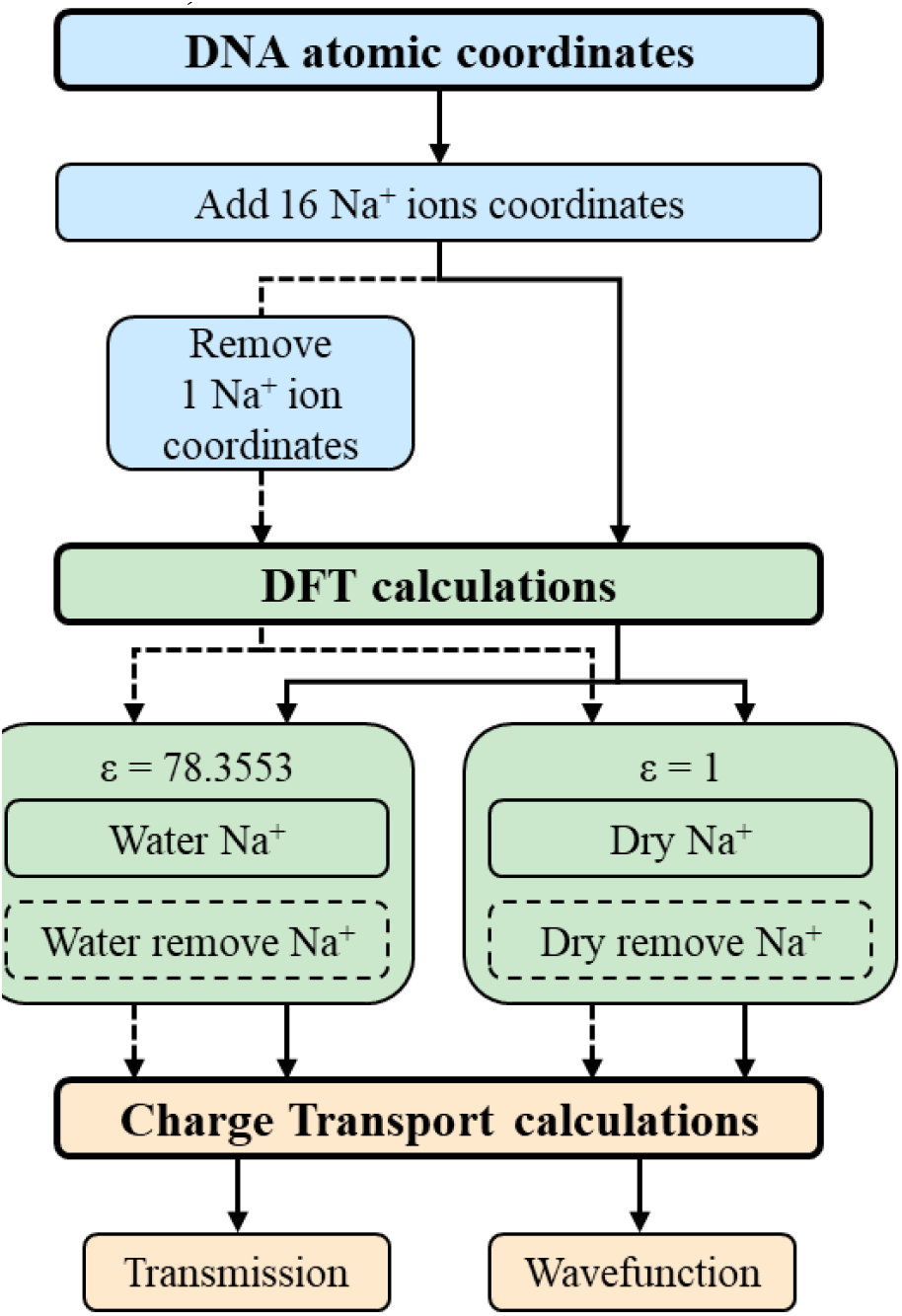
The flow chart of simulation procedures.

### A. DFT calculations

We first obtain the atomic coordinates of the double-stranded B-DNA using the Nucleic Acid Builder [42]. Then, we add counterions along the backbone using the approach in Qi et al. [43]. The minimization of Na^+^ ions is performed with a single strand of Bconformation DNA consisting of three bases and the phosphate backbone while the DNA is held fixed. The relative dielectric constant of unity (dry) was used. Qi et al. found that the Sodium atom should be at a location of 2~3 Å away from the phosphate group (see Section III of Supplemental Material for the precise coordinates used). We followed this with DFT calculations to find the Fock and Overlap matrices of the DNA strand with counterions. As we are probing the role of the solvent dielectric in this work, we keep the coordinates of the DNA and counterions fixed. The only difference between dry (dehydrated) and water (hydrated) DNA is the value of the dielectric constant in the DFT calculations. While energy minimized coordinates for atoms in dry DNA would be more appropriate, force fields for the dehydrated condition are not available, and so in this computational study, we use the same coordinates in both cases. In a solvent environment, we expect the location of the counterion to be further away from the phosphate.

In calculating the impact of the solvent, abinitio studies have concluded that the polarizability of the solvent is the essential factor affecting ionization potential [44–46]. Studies show that the polarizable continuum model (PCM) captures the screening effect of the solvent without the need to include explicit water molecules [46]. Therefore, to account for the water solvent, we use the PCM. The DFT calculations are carried out with the B3LYP/6-31G(d,p) basis set [47] with one counterion at every phosphate backbone, with the total charge of the system being zero. We choose this basis set based on a balance between calculation accuracy and reasonable computational cost. We verified that our results are consistent with different basis sets: B3LYP/6-311G(d,p), B3LYP/cc-pVDZ, and B3LYP/cc-pVTZ. Please refer to Section I of Supplemental Material for detailed discussions. Note that the terminal bases at the 5’ end do not include the phosphate groups. After reaching convergence, the Fock and overlap matrices obtained from the DFT calculations are used in the transport calculations, which is the third and final step. To understand the role of counterions in influencing transport, we randomly select one of the 16 Na^+^ ions and remove its coordinate right before DFT calculation. We refer to these calculations as the “Removed Na^+^” case, and the total charge of the system is set to -1. The simulation procedure is kept identical for all cases. Thus, overall, we investigate four different cases: *Water Na*^*+*^ (ε = 78.3553, with 16 Na^+^ ions), *Water Removed Na*^*+*^ (ε = 78.3553, with 15 Na^+^ ions), *Dry Na*^*+*^ (ε = 1, with 16 Na^+^ ions), and *Dry Removed Na*^*+*^ (ε = 1, with 15 Na^+^ ions).

### B. Charge transport calculations

Phase coherent transmission of electrons from one contact to the other through the DNA involves using the Hamiltonian from the DFT calculations discussed in Section II A (called the coherent case). In the coherent case, the quantum mechanical phase of the electron evolves as per the single particle Schrodinger equation, and the electron does not feel the influence of the other degrees of freedom such as lattice vibrations and solvent environment. We know from prior work that the coherent case yields very low values of the transmission compared to experiments. Decoherence, which represents the interaction of the effective single particle Hamiltonian with other degrees of freedom, helps us move closer to explaining a set of experiments [43]. Thus, here, we present results for the decoherent case. For the comparison between coherent and decoherent cases, we refer readers to Section II of Supplemental Material.

The charge transport calculations are carried out using the Green’s function method by closely following the method used in references [43] and [48]. To model decoherence, we used decoherence probes at each atom in the system [48]. Our primary constraint is setting the net current at each probe equal to zero. The electrical contacts are made at the cytosine bases in the 3’-end and 5’-end to mimic an experimental configuration (see FIG. 2).

**FIG. 2.**
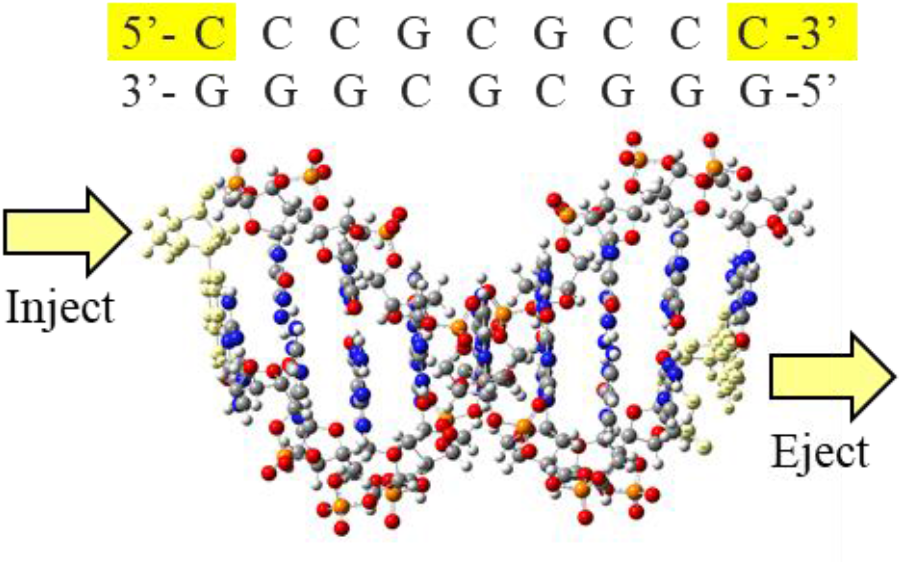
The sequence and the atomic structure of the B-DNA strand. The yellow arrows represent the contact / open boundary conditions. The yellow highlight atoms on the two ends are where the contact self-energies are applied.

We obtain the Fock (*H*_0_) and overlap (*S*_0_) matrices from the DFT calculations (Section II A), we set the dielectric constant to be 78.3553 and 1.0 for wet and dry conditions, respectively via the PCM model. Using Löwdin transformation, the non-orthogonal basis set Fock matrix *H*_0_ is converted to an orthogonal basis set Hamiltonian *H*:

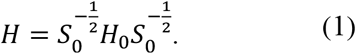

The diagonal elements of *H* represent the energy levels at each atomic orbital of the system. The off-diagonal elements of *H* represent the coupling between the different atomic orbitals. With energy levels and coupling in place, we used the Green’s function method; in particular, the retarded Green’s function (*G*^*r*^), is calculated using:

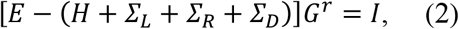

where *E* is the energy and *H* is the Hamiltonian. Σ_*L*(*R*)_ is the left (right) contact retarded self-energy, which represents the coupling strength of the contacts to the DNA molecule. The selfenergy Σ_*D*_ is due to the decoherence probes. Using the wide-band limit approximation [49], we assume an energy-independent self-energy, which is defined as

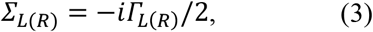

where *i* is the imaginary unit, and Γ_*L*(*R*)_ the coupling strength between the DNA and the left (right) contact.

The self-energy due to the phase breaking probe (Σ_*D*_) represents the influence of these probes on the DNA. They are defined in a similar manner:

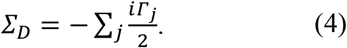

The summation over j on the right-hand side is over the probes. 𝛤_*j*_ represents the coupling strength between the probe and the DNA is taken as an energy-independent parameter.

We set the left (right) contact scattering rate 𝛤_L_(𝛤_R_) to 100 meV and the decoherence scattering rate to 10 meV, which are within the acceptable range [43,50]. The temperature is assumed to be 298K. The current at the *i*^*t*h^ probe is defined as

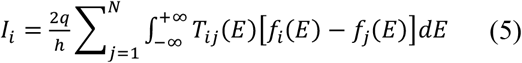

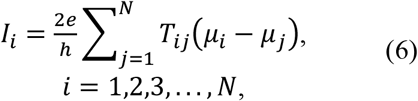

where *T*_*ij*_ = *Γ*_*i*_*G*^*r*^*Γ*_*j*_*G*^*a*^ is the transmission probability between the *i*^*t*h^ and *j*^*t*h^ probes, *G*^*a*^ = (*G*^*r*^)^†^ is the advanced Green’s function, 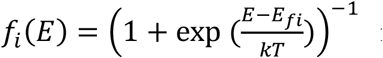 is the Fermi distribution. *N* is the total number of probes in the system (including the contact atoms) and *N*_*contacts*_ is the total number of probes on contact atoms.

To ensure current continuity, the current at each probe with respect to energy is set to zero. In our calculations, we applied the decoherence probes at each atom, thus, we have number of probes *N*_*b*_ = *N* − *N*_*contacts*_ This condition yields *N*_*b*_ independent equations that help derive the following relation [51].

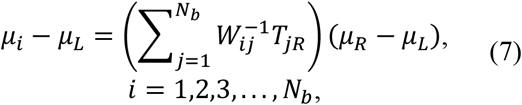

where 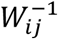 is the inverse of *W*_*ij*_ = (1 − *R*_*ii*_)δ_*ij*_ − *T*_*ij*_(1 − δ_*ij*_), and *R*_*ii*_ is the reflection probability at probe *i*, and is given by 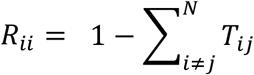. Further, since the current at the *i*≠*j* left and right contacts is not zero, we can write the equation for current as

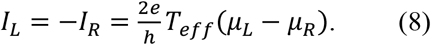

Comparing Eq. (6) to Eq. (8), we obtain the effective transmission:

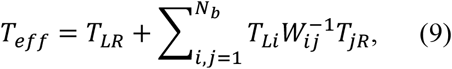

where *T*_*LR*_ is the coherent transmission from the left to the right contact, and the second term is the decoherent transmission component.

## III. RESULTS AND DISCUSSION

To study the role of static counterions (Na^+^) and solvent in affecting the charge transport, we start by investigating *Water Na*^*+*^ and *Dry Na*^*+*^ cases. We use the PCM with the dielectric constant (ε) to model the solvent. We first observe that the bandgap significantly depends on the dielectric constant. For *Dry Na*^*+*^ case, the bandgap is 0.77 eV, much smaller than the value of 4.03 eV for the *Water Na*^*+*^ case (see the first row of TABLE I).

**TABLE I:** HOMO band, LUMO band, and Bandgap comparisons between four different cases.

| | | Water ($\epsilon = 78.3553$) | | Dry ($\epsilon = 1$) | |
| --- | --- | --- | --- | --- | --- |
| | | Removed $\text{Na}^+$ | $\text{Na}^+$ | Removed $\text{Na}^+$ | $\text{Na}^+$ |
| Bandgap (eV) |  | 4.0795 | 4.0333 | 0.4560 | 0.7714 |
| HOMO<br>Band | HOMO (eV) | -5.0708 | -5.0771 | -2.1045 | -3.4365 |
|  | Location | Guanine | Guanine | Guanine | Guanine |
|  | Levels | 9 | 9 | 10 | 9 |
|  | Band width (eV) | 0.3941 | 0.3927 | 1.3323 | 1.0344 |
| | Level density ($\text{eV}^{-1}$) | 22.8368 | 22.9183 | 7.5058 | 8.7007 |
| LUMO<br>Band | LUMO (eV) | -0.9913 | -1.0438 | -1.6485 | -2.6651 |
| | Location | Cytosine | Cytosine | $\text{Na}^+$ | $\text{Na}^+$ |
|  | Levels | 9 | 9 | 15 | 16 |
|  | Band width (eV) | 0.2204 | 0.2468 | 1.3353 | 1.2952 |
| | Level density ($\text{eV}^{-1}$) | 40.8348 | 36.4668 | 11.2334 | 12.3533 |

The transmission is a measure of electron flow between the left and right contacts through the DNA (FIG. 2). As discussed in the Methods section, the electrons interact with the decoherence probes as they flow through the DNA. It has been previously shown that the conductance of DNA molecules can be altered by conformation and solvent environments [15,35–37]. However, their individual effect in determining the conductance is hard to distinguish in prior modeling studies. Our goal is to systematically understand how counterions and solvent dielectric individually influence the transmission. Therefore, we keep the DNA coordinates or geometry fixed throughout our calculations. And we compare our results for transmission obtained from the same DNA structure with counterions in the dry and water environment.

FIG. 3 shows that the transmission of *Water Na*^*+*^ case at the HOMO and LUMO bands is primarily through the atoms of the DNA bases rather than Na^+^ ions. This is further enunciated by plotting the wavefunctions of the highest nine HOMO and the lowest nine LUMO energy levels in FIG. 3 (b) and (c). The wavefunctions at these energy levels all lie on the Guanines (HOMO band) and Cytosines (LUMO band). A previous computational study [52] used structures with Na^+^ ions placed further away than in our calculations (2~3 Å from the phosphate group), and they found that the HOMO and LUMO levels are determined by the DNA bases, similar to our results. In FIG. 3 (a), we note that the band of energies around 0 eV is due to electron transport along the Na^+^ ions since the wavefunctions of these energy levels are primarily localized on Na^+^ ions.

**FIG. 3.**
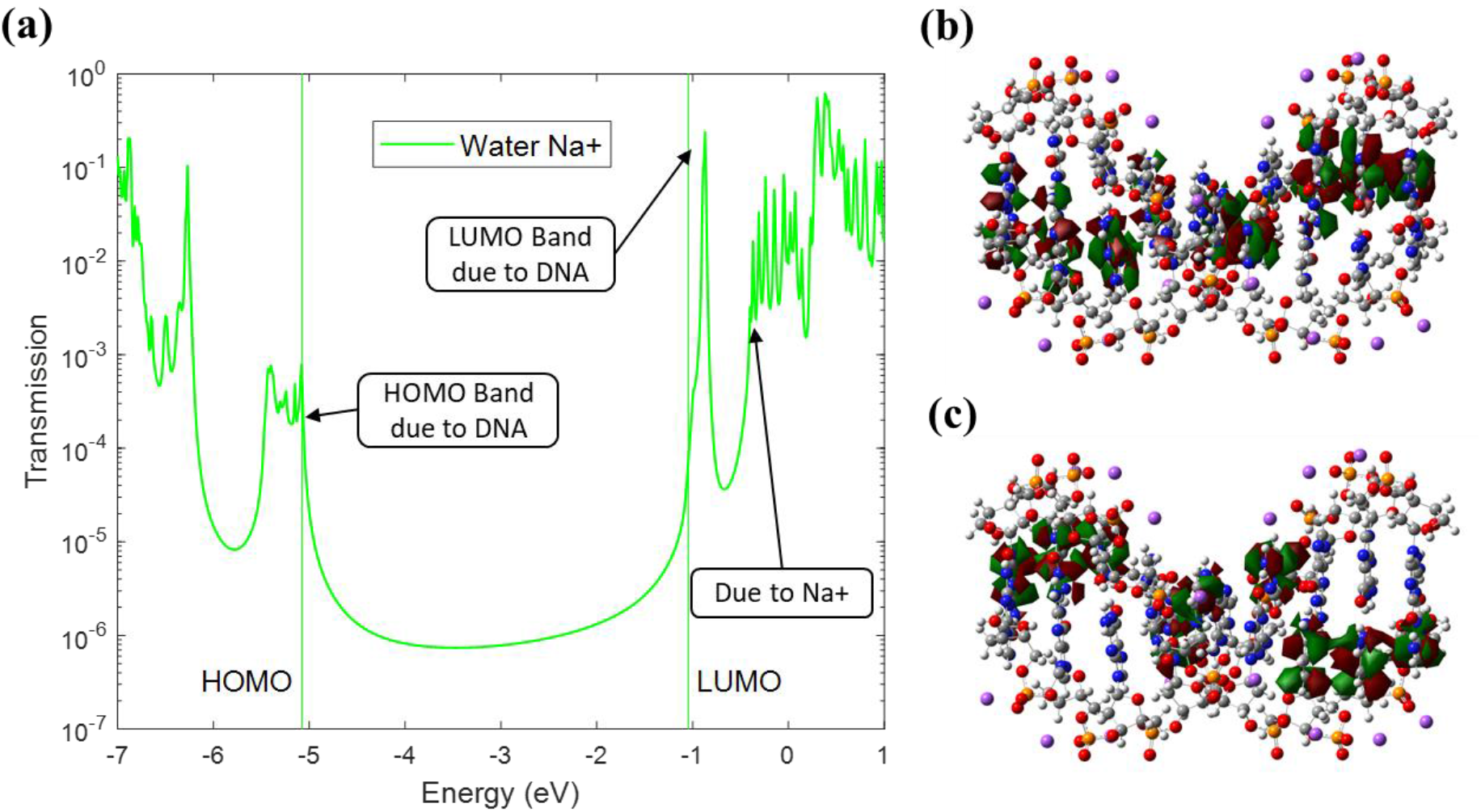
(a) Decoherent transmission of DNA with Na^+^ ions in water environment (*Water Na*^*+*^). The arrowedboxes indicate the localization of several energy levels based on wavefunction plots. (b) The wavefunctions of the highest nine HOMO energy levels (HOMO band) are localized on Guanine bases. (c) The wavefunctions of the lowest nine LUMO energy levels (LUMO band) are localized on Cytosine bases.

For *Dry Na*^*+*^ case, we also observe that the transmission at the HOMO band is through the DNA molecule (see FIG. 4). FIG. 4 (b) shows that the highest nine HOMO band wavefunctions lie on the Guanines. In contrast, the transmission at the LUMO band present around energies of -2 eV is due to Na^+^ ions. The wavefunctions corresponding to the lowest sixteen LUMO energies all lie on the Na^+^ ions, as shown in FIG. 4 (c). The transmission resonances at the unoccupied orbitals around−0.8 *eV* is due to transport through both the DNA molecule and the Na^+^ ions because the wavefunctions at these energies are localized on either Cytosines or Na^+^ ions.

**FIG. 4.**
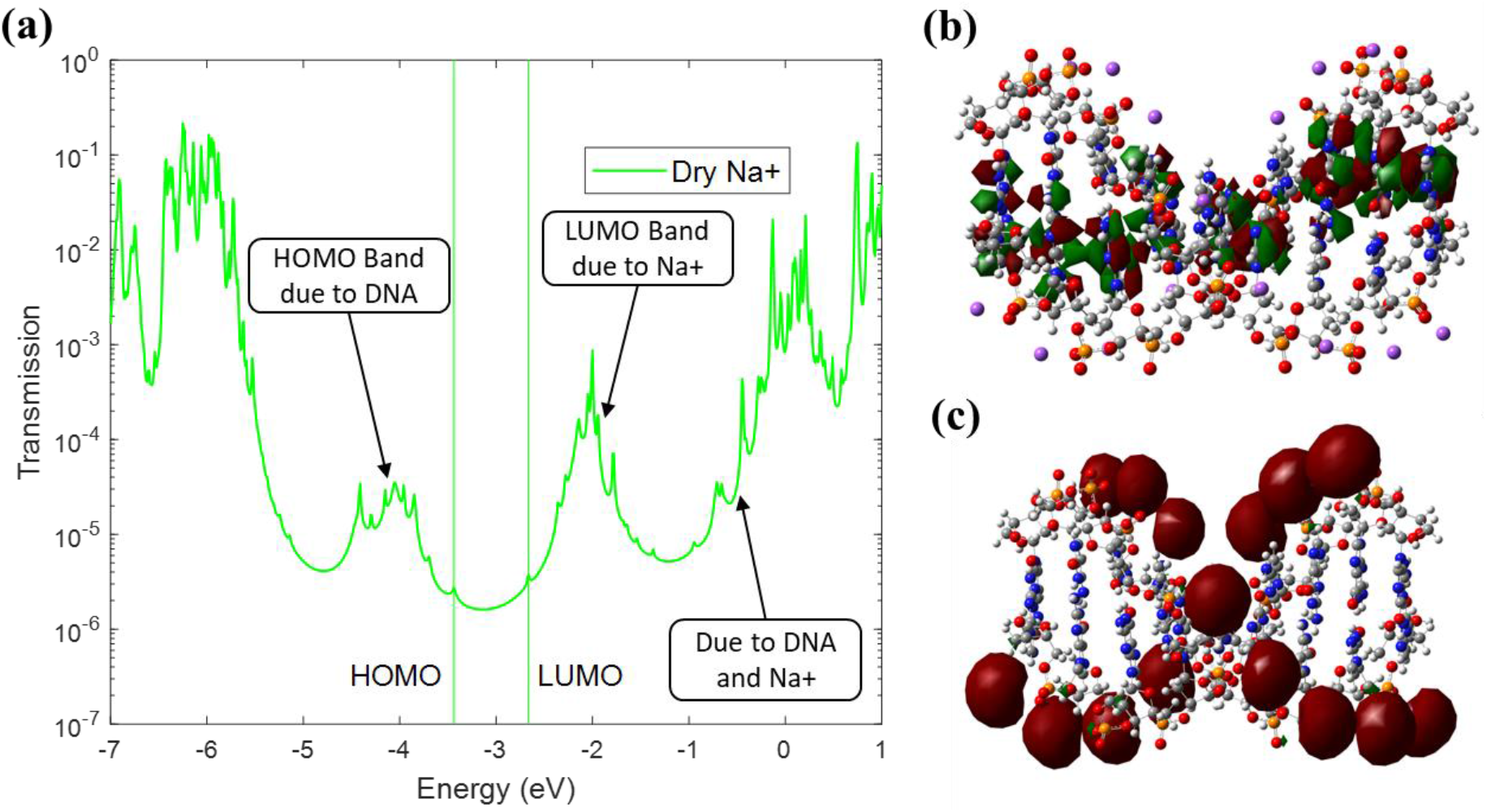
(a) Decoherent transmission of DNA with Na^+^ ions in a dry environment (*Dry Na*^*+*^). The arrowed-boxes indicate the localization of several energy levels based on wavefunction plots. (b) The wavefunctions of the highest nine HOMO energy levels (HOMO band) are localized on Guanine bases. (c) The wavefunctions of the lowest sixteen LUMO energy levels (LUMO band) are localized on Na^+^ ions. The energies above the LUMO band (−1 eV onwards) consist of energy levels on the Cytosine and Na^+^ ions (not shown here). See Section II of Supplemental Material for coherent results, which lend further support to these observations.

Although the actual HOMO energy level is essential, the energy separation between molecular orbitals and their spatial distribution is also critical for the electrons to hop from one energy level to another, which leads to better transmission. For easier comparison, we define the number of energy levels divided by the band width as *level density*. We observe that the *level density* depends on the dielectric constant from the results summarized in TABLE I. In general, *Water* cases have higher level density than *Dry* cases, thus larger transmission. Similarly, LUMO bands have higher level density than HOMO bands, thus larger transmission. Although the transmission is not linearly proportional to level density, their correlation is not neglectable.

To further investigate the role of counterions, we performed a simulation by randomly removing one of the 16 Na^+^ counterions from the DNA model and setting the system’s total charge to negative one. For instance, by removing the 7^th^ Na^+^ ion from the *Water Na*^*+*^ case, no significant difference in transmission occurs at the HOMO and LUMO energies, as seen in FIG. 5 (a). Only the transmission peaks at energies above the LUMO band become smaller. This characteristic, besides the wavefunction, provides evidence that the band of energy levels around 0 eV is formed by Na^+^ ions. The level densities of both HOMO and LUMO bands stay relatively constant with or without removing one of the 16 Na^+^ ions.

**FIG. 5.**
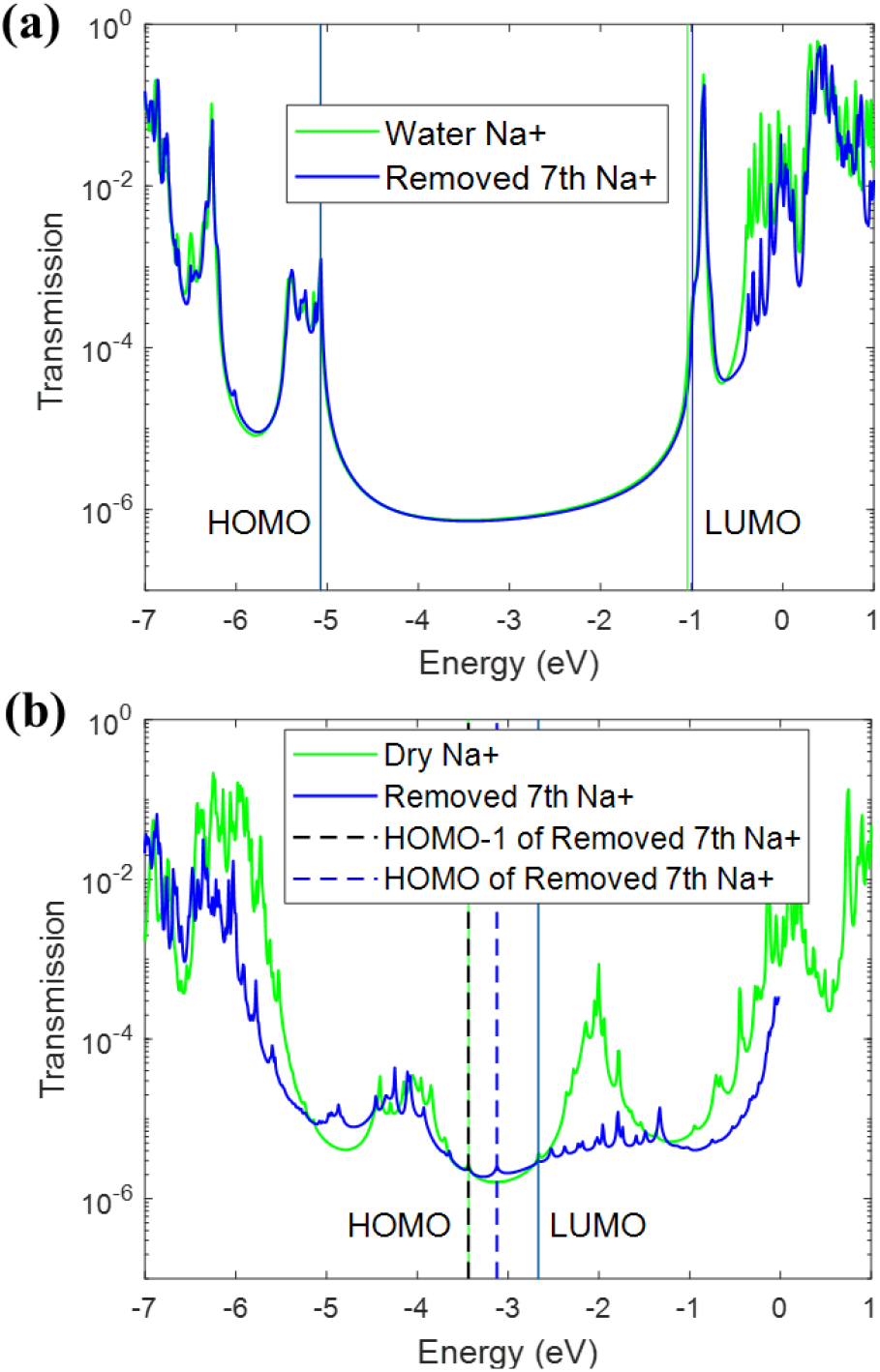
Comparison of the transmission plots with (Green) and without (blue) the 7^th^ Na^+^ ion removed. All other counterions are present. (a) No significant difference is seen with the water solvent. (b) A significant difference is observed for *Dry* cases at both the HOMO and LUMO bands. For subplot (b), we have shifted the energy to align the LUMO level of both molecules for easy comparison.

In comparison, removing the 7^th^ Na^+^ ion from *Dry Na*^*+*^ case lowers the transmission of the LUMO band significantly in FIG. 5 (b). Missing one Na^+^ ion breaks the transport pathway of electrons, which is clearly built on 16 Na^+^ ions. While the LUMO band width stays relatively constant, the level density decreases by about 1.1 eV^-1^ after losing one energy level. Therefore, in FIG. 5 (b), the LUMO level of both dry cases are aligned for easy comparison. Meanwhile, we noticed HOMO-1 of *Dry Removed Na*^*+*^ case coincides with the HOMO of *Dry Na*^*+*^ case. A higher occupied energy level appears in *Dry Removed Na*^*+*^ case beyond the original HOMO band of *Dry Na*^*+*^ case. The location of this new HOMO energy level further decreases the bandgap to 0.46eV. One extra energy level in HOMO band causes its band width to extend by about 0.3 eV and its level density to decrease by about 1.2 eV^-1^. As a result, the average transmission value across the HOMO band is slightly lower than *Dry Na*^*+*^ case.

To understand the underlying reasons for the above observations on the role of the solvent and counterions on transport properties, we analyze the width of the HOMO and LUMO bands, the onsite potential at the bases, and the hopping strength between neighboring bases. For this, we first arrange the Hamiltonian *H* (from Eq. (1)) in the order of DNA bases,

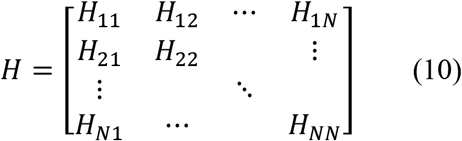

where *N* is the number of bases (*N* = 18). The diagonal subblocks *H*_*ii*_ correspond to the Hamiltonian of base *i*, and the off-diagonal subblock *H*_*ij*_ represents the coupling between bases *i* and *j*. Then we perform the following transform to diagonalize all diagonal subblocks of *H*,

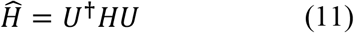

where *U* is a block diagonal matrix. To obtain *U*, we calculate the eigenvectors of each subblock of *H*, then construct the entire *U* matrix,

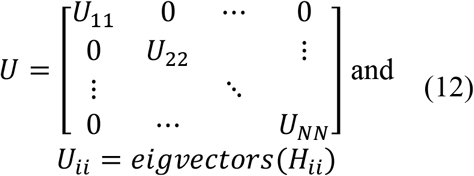

The resulting Hamiltonian *Ĥ* is similar in form to Eq. (10). The diagonal subblocks of *Ĥ* correspond to the energy levels of each DNA base, and the off-diagonal subblocks represent the hopping strength between energy levels on two different bases. Finally, in our discussion below, we restrict ourselves to one energy level (such as HOMO or LUMO).

The HOMO band of water case lies from -5.47eV to -5.08eV, corresponding to a bandwidth 0.39eV (see TABLE I). In comparison, the HOMO level density of *Dry* case is about 2.6 times smaller. The underlying reason for this is the low dielectric constant in the Dry case. It results in a larger separation (smaller level density) between the onsite energy levels at the bases corresponding to the HOMO band. On the other hand, the high dielectric constant of water makes the onsite energies corresponding to the HOMO band energetically closer (larger level density).

The onsite potential of the bases corresponding to the HOMO band in the water and dry cases are shown in FIG. 6. Although the hopping terms are similar in the two cases (especially coupling between G-G neighbor bases), the onsite potentials of the water case are more uniform and closer together. This behavior can also be seen by comparing their coefficient of variation which are −0.0581 (water case) vs. −0.1811 (dry case). The HOMO level density of the water case is 22.92 eV^-1^, which is about 2.6 times larger than the dry case (8.70 eV^-1^). As a result, the average transmission value across the HOMO band is approximately one to two orders of magnitude larger in the water case as opposed to the dry case. Therefore, the electrons travel through the DNA more efficiently in the water case.

**FIG. 6.**
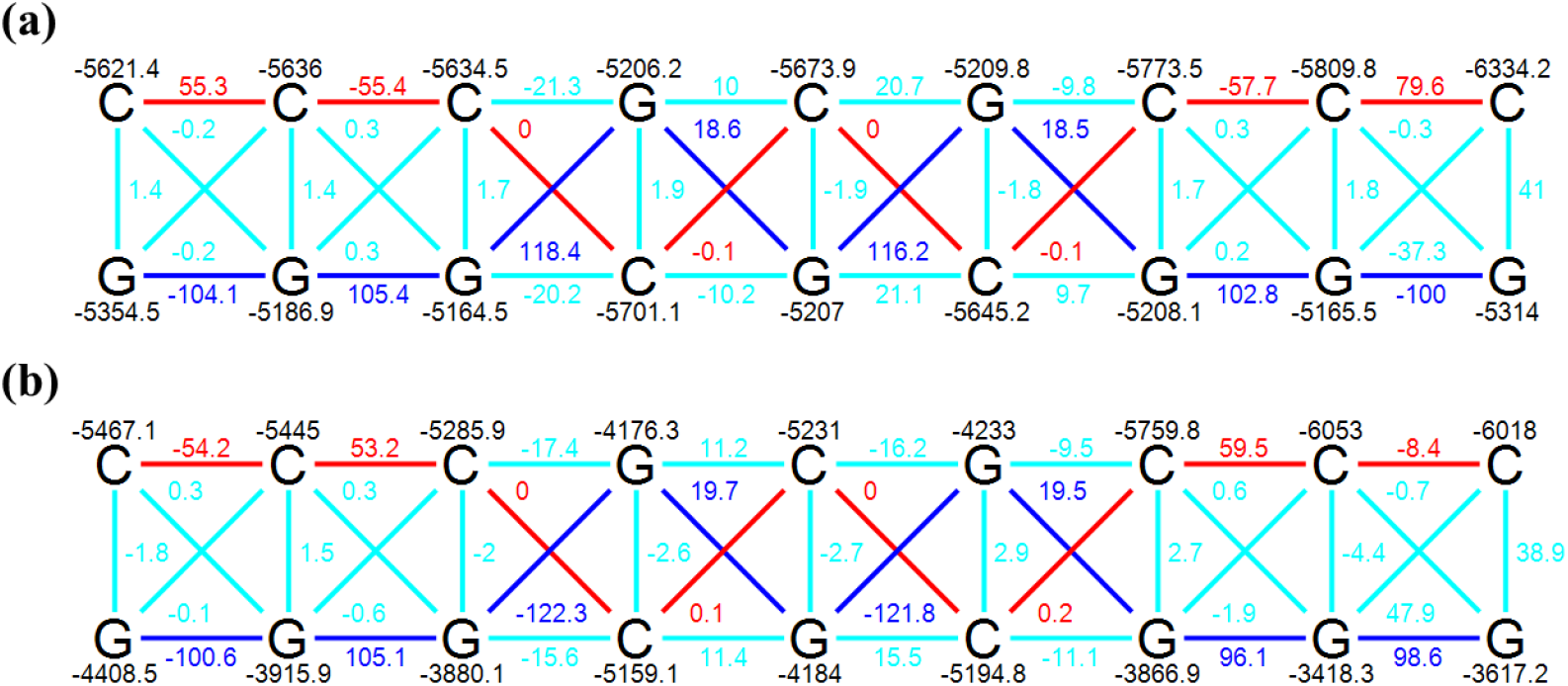
The hopping parameters between neighboring bases at (a) the highest 18 HOMO levels of *Water Na*^*+*^ case and (b) the highest 18 HOMO levels of *Dry Na*^*+*^ case. Units are in meV. For (a) and (b), the coefficient of variation of the onsite potentials is −0.0581 and −0.1811, respectively.

The onsite potential of the bases corresponding to the LUMO band in the water and dry cases are shown in FIG. 7. Unlike all other bands, the LUMO band of *Dry Na*^*+*^ case comprises 16 energy levels localized on the 16 Na^+^ ions instead of DNA molecule. The onsite potentials of the Na^+^ ions in the LUMO band range in energies from -2.43eV to -1.45eV. The hopping terms between Na^+^ ions in the LUMO band are primarily in the range of 50 to 70 meV.

**FIG. 7.**
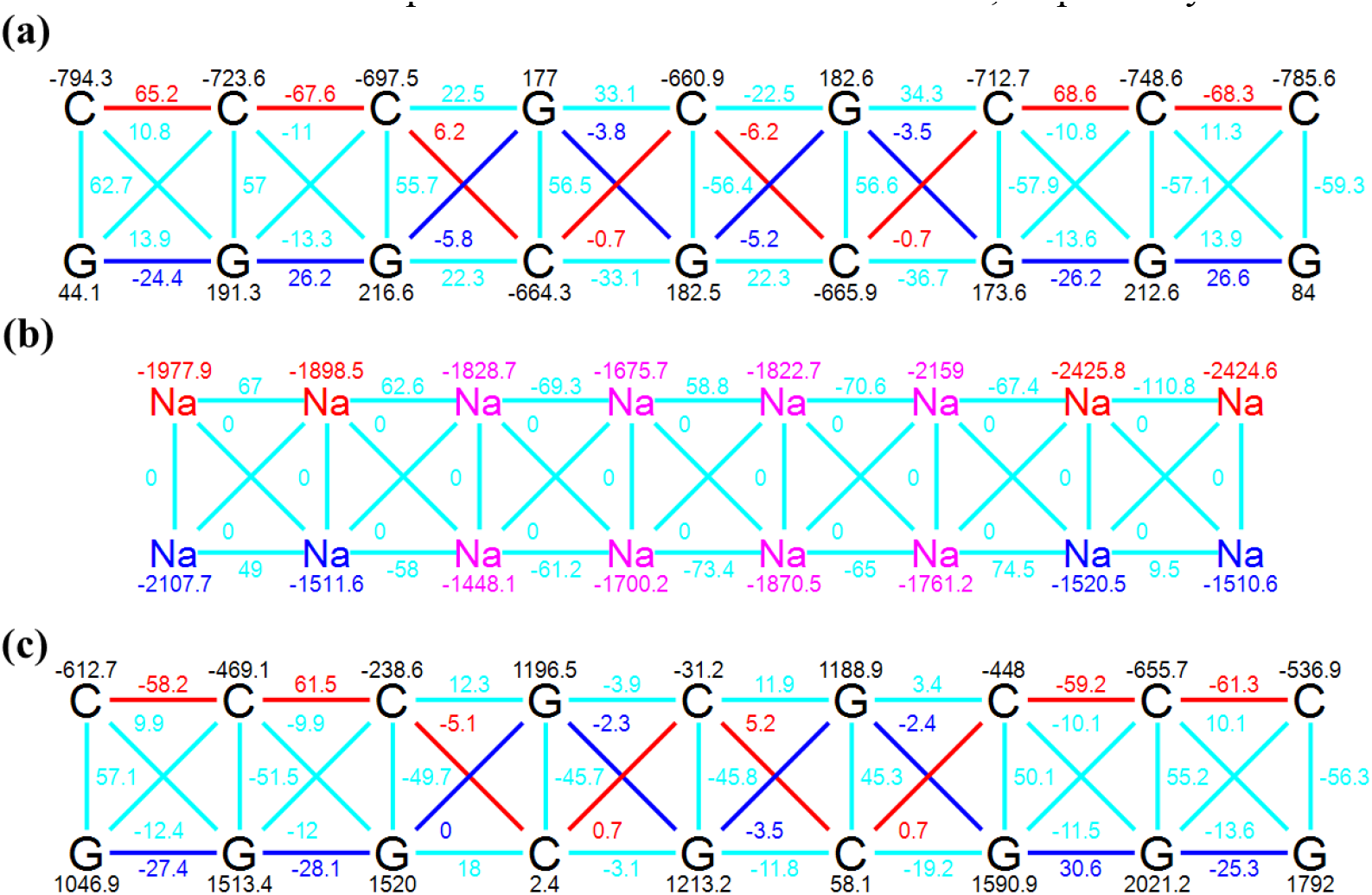
The hopping parameters between neighboring bases at (a) the lowest 18 LUMO levels of *Water Na*^*+*^ case, (b) the lowest 16 LUMO levels of *Dry Na*^*+*^ case, and (c) the even higher LUMO levels (LUMO + 16 ~ LUMO + 33) of *Dry Na*^*+*^ case. Units are in meV.

## IV. CONCLUSION

It has been challenging to obtain clear trends in the underlying physics of DNA conduction due to the complexity of environmental conditions. This situation has motivated us to computationally study an important aspect encountered - the effect of the solvent and counterions for a static configuration of atoms. We consider a nine-base-pair double-stranded B-DNA and study the role of counterion arrangement and solvent dielectric constant to determine if there are clear trends in the underlying physics. We use the PCM model for the solvent and consider the dry and fully hydrated environments. By performing calculations on six different DNA sequences, we emphasize the generalizability of the results (additional results are presented in Section III of Supplemental Material).

Depending on the dielectric constant of the surrounding medium, Na^+^ ion is found to impact the charge transport properties of the DNA significantly. From the molecular energy level perspective, Na^+^ ions add unoccupied energy levels in the bandgap of the DNA in the dry case. On the other hand, the water case adds unoccupied energy levels that have higher energy than the LUMO, which is primarily located on the cytosine bases. Because of the high dielectric constant of water, the interaction between DNA and Na^+^ ions, is effectively screened. In addition, from the charge transport perspective, the transmission is at least two orders of magnitude larger at HOMO and LUMO regions of the DNA in the water case than in the dry case. The observed narrower spread of onsite potentials (at HOMO and LUMO bands) with a water environment supports higher transmission.

In summary, our simulation results demonstrate that it is essential to consider counterions as an individual factor when analyzing the DNA conductance experiments done in the dry case but not necessarily in the water solvent. The higher the dielectric constant, the higher the charge screening effect, thus lowering the coupling between Na^+^ ions and DNA molecules. As the presence of Na^+^ ions added energy levels within the bandgap of the DNA in dehydrated condition (the dry case), this can further be relevant to utilizing DNA in nanoelectronics applications.

## ACKNOWLEDGMENTS

We acknowledge National Science Foundation Grant Numbers 1807391/1807555 (SemiSynBio Program) and 2036865 (Future of Manufacturing) for support.

