## Supplemental Material for "A Computational Study of the Role of Counterions and Solvent Dielectric in Determining the Conductance of B-DNA"

### I. Larger Basis Set Results

In the context of the level of theory for the density functional theory (DFT) calculations, the B3LYP/6-31G(d,p) has been used to calculate the ionization potential of nucleobases extensively. These calculations find the B3LYP/6-31G(d,p) to yield the correct trend in ionization potential as experiments but with an offset in values of about 300 meV. Other more expensive methods, such as CCSD, MP2, and cc-pVTZ, may give a higher accuracy for the ionization potential, but what makes B3LYP/6-31G(d,p) a method of choice is a balance between calculation accuracy and reasonable computational cost. Since our system under study comprises a relatively large molecular structure (~700 atoms), we need to use the entire Hamiltonian and Overlap matrices to calculate the transport properties.

To demonstrate that our results are more or less basis-set-independent, we performed the same simulation procedure as the one discussed in Section II of main manuscript on the sequence of 5'-CCCGCGCCC-3' with three different basis sets for DFT calculation: (a) B3LYP/6-311G(d,p), (b) B3LYP/cc-pVDZ, and (c) B3LYP/cc-pVTZ. Please note that B3LYP/6-311G(d,p) and B3LYP/cc-pVTZ are larger basis sets than B3LYP/6-31G(d,p). To evaluate the robustness, we focused on B3LYP/cc-pVTZ as a benchmark since it has been shown to yield more accurate results but with an increase in the computational time. The transmission and wave function plots calculated using B3LYP/cc-pVTZ are shown in the following figures. A comparison between calculations (Fig. 3 vs. FIG. S1 and Fig. 4 vs. FIG. S2) indicates that using B3LYP/cc-pVTZ basis set produces results consistent with the B3LYP/6-31G(d,p) basis set. Analysis of the LUMO orbitals under dehydrated condition (see FIG. S2) shows that orbitals are still localized on $Na^+$ ions when the cc-pVTZ basis set is used. Similarly, in the water solvent (as shown in FIG. S1), HOMO orbitals continue to be localized on Guanine bases. In addition, the transmission and wave function plots, calculated using B3LYP/6-311G(d,p) and B3LYP/cc-pVDZ, show the same behavior (see Fig. S3, S4, S5, S6). Therefore, we anticipate the B3LYP/6-31G(d,p) basis set provides a balance between accuracy and computational cost and the conclusions drawn in the paper are accurate enough to be considered as robust and independent of the choice of basis sets.

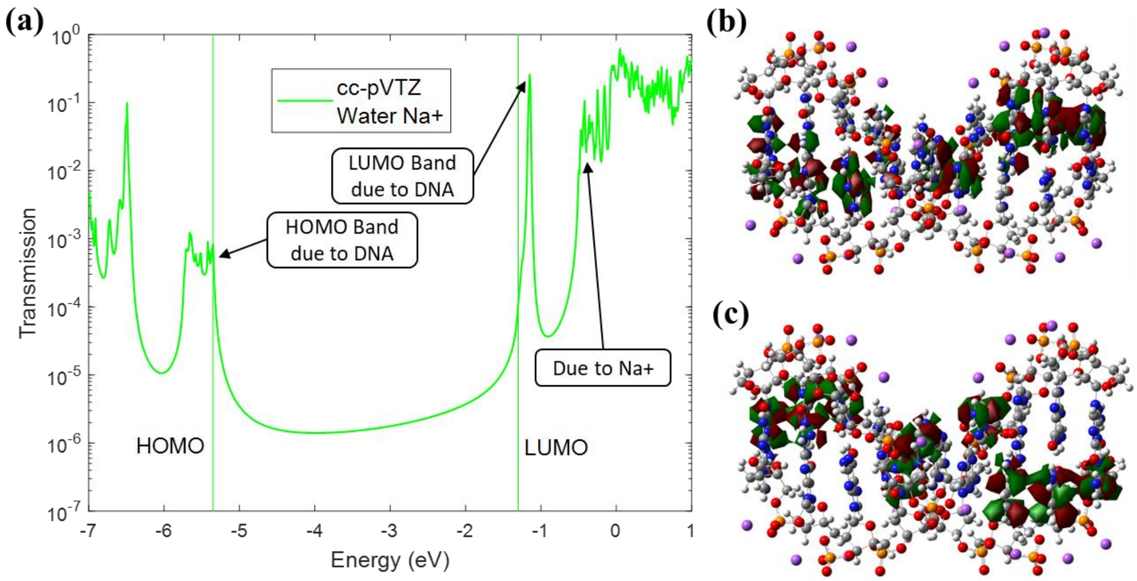


FIG. S1. DFT calculations with B3LYP/cc-pVTZ basis set for (a) Decoherent transmission of DNA with $Na^+$ ions in water environment (*Water* $Na^+$). The arrowed-boxes indicate the localization of several energy levels based on wavefunction plots. (b) The wavefunctions of the highest nine HOMO energy levels (HOMO band) are localized on Guanine bases. (c) The wavefunctions of the lowest nine LUMO energy levels (LUMO band) are localized on Cytosine bases.

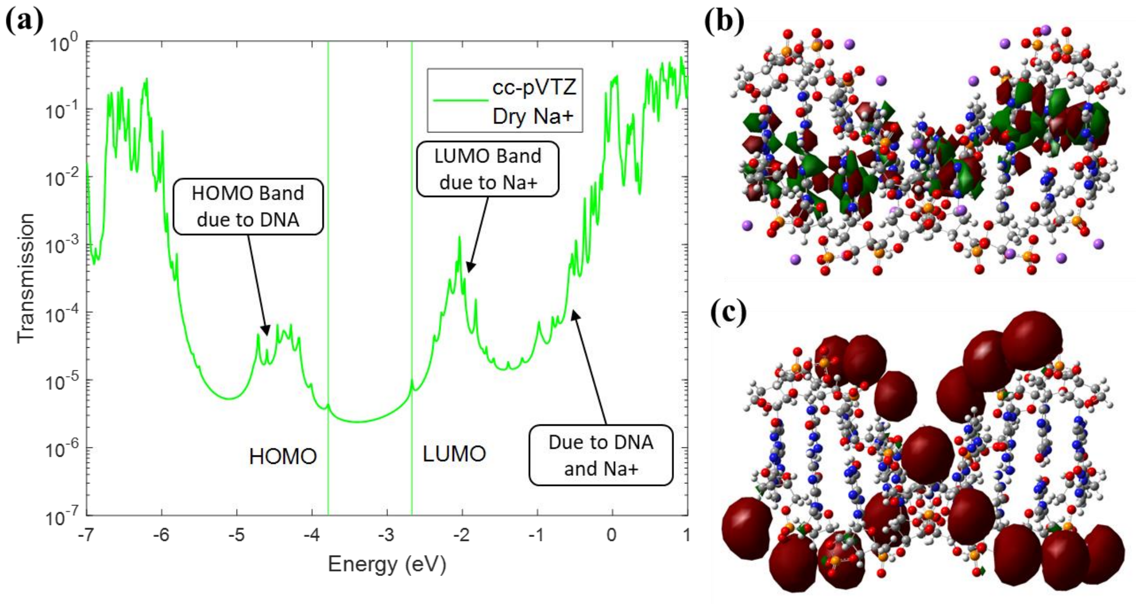


FIG. S2. DFT calculations with B3LYP/cc-pVTZ basis set for (a) Decoherent transmission of DNA with $Na^+$ ions in a dry environment (*Dry* $Na^+$). (b) The wavefunctions of the highest nine HOMO energy levels (HOMO band) are localized on Guanine bases. (c) The wavefunctions of the lowest sixteen LUMO energy levels (LUMO band) are localized on $Na^+$ ions.

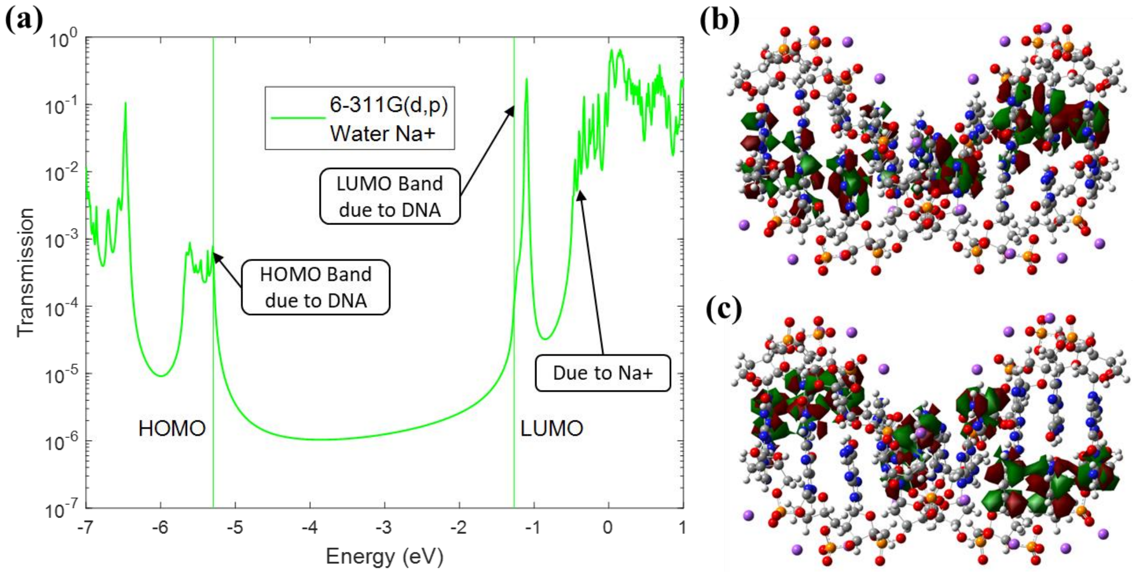


FIG. S3. DFT calculations with B3LYP/6-311G(d,p) basis set for (a) Decoherent transmission of DNA with $Na^+$ ions in a water environment (*Water* $Na^+$). (b) The wavefunctions of the highest nine HOMO energy levels (HOMO band) are localized on Guanine bases. (c) The wavefunctions of the lowest nine LUMO energy levels (LUMO band) are localized on Cytosine bases.

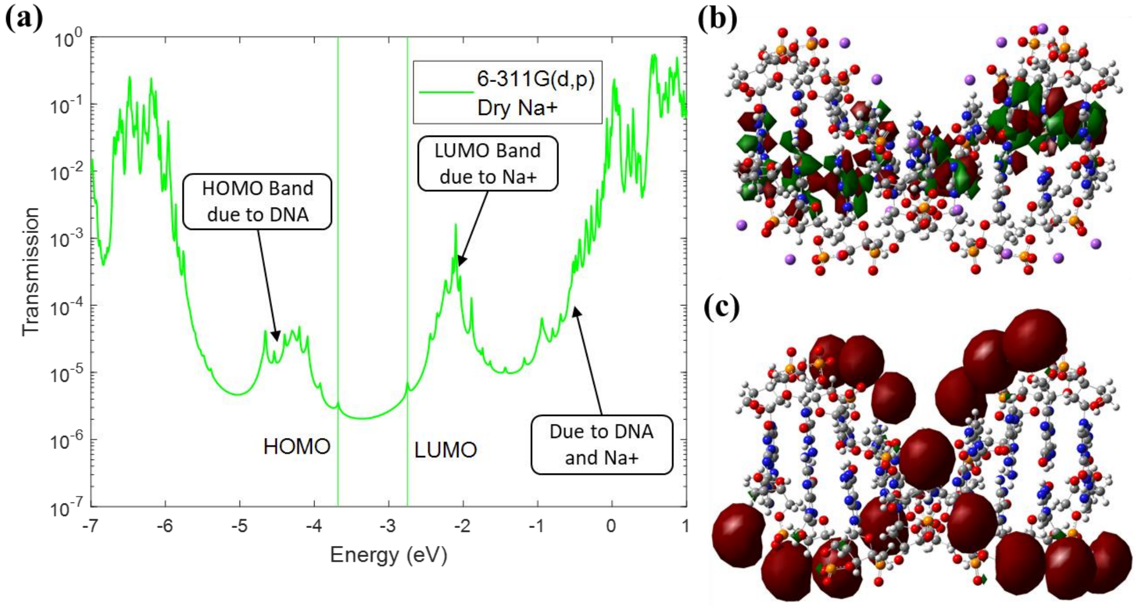


FIG. S4. DFT calculations with B3LYP/6-311G(d,p) basis set for (a) Decoherent transmission of DNA with $Na^+$ ions in a dry environment (*Dry* $Na^+$). (b) The wavefunctions of the highest nine HOMO energy levels (HOMO band) are localized on Guanine bases. (c) The wavefunctions of the lowest sixteen LUMO energy levels (LUMO band) are localized on $Na^+$ ions.

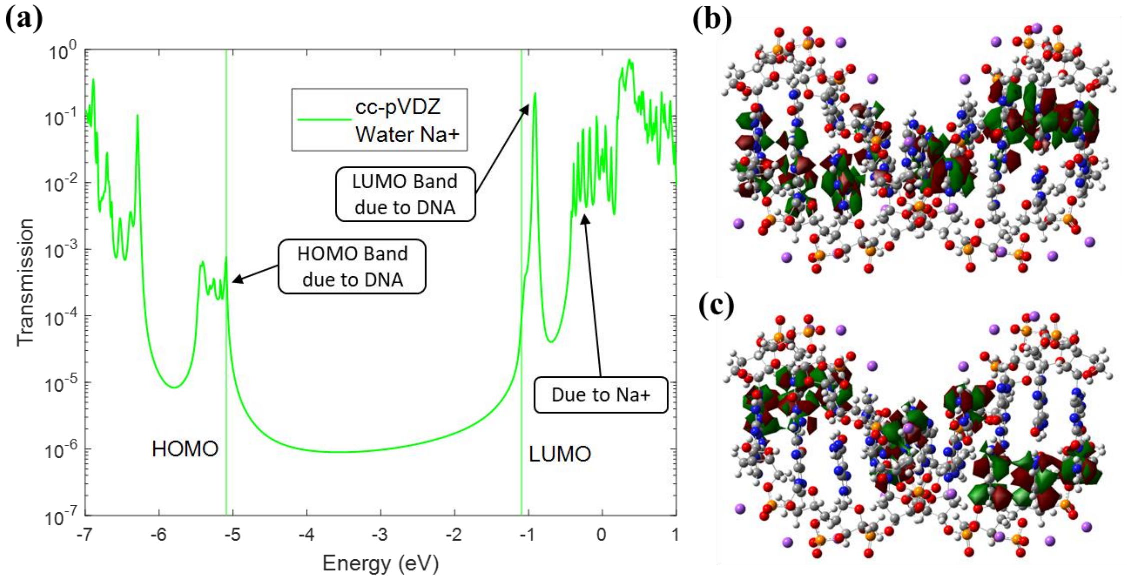


FIG. S5. DFT calculations with B3LYP/cc-pVDZ basis set for (a) Decoherent transmission of DNA with $Na^+$ ions in a water environment (*Water* $Na^+$). (b) The wavefunctions of the highest nine HOMO energy levels (HOMO band) are localized on Guanine bases. (c) The wavefunctions of the lowest nine LUMO energy levels (LUMO band) are localized on Cytosine bases.

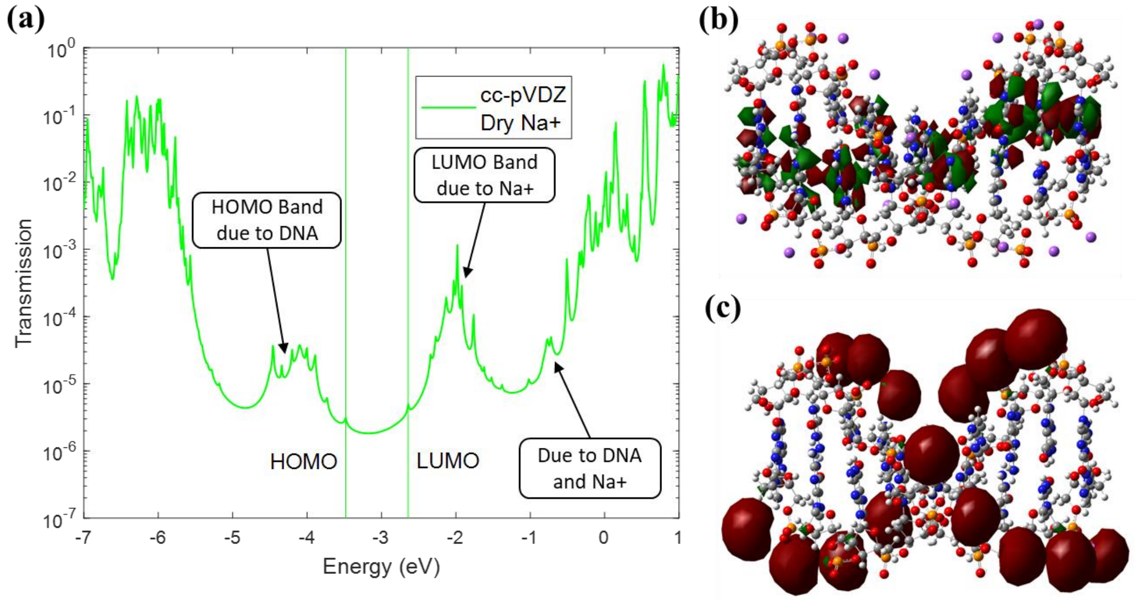


FIG. S6. DFT calculations with B3LYP/cc-pVDZ basis set for (a) Decoherent transmission of DNA with $Na^+$ ions in a dry environment (*Dry* $Na^+$). (b) The wavefunctions of the highest nine HOMO energy levels (HOMO band) are localized on Guanine bases. (c) The wavefunctions of the lowest sixteen LUMO energy levels (LUMO band) are localized on $Na^+$ ions.

### II. Coherent Transmission Results

We performed coherent transmission calculations along with decoherent ones which are extensively discussed in the main manuscript. As discussed in the main manuscript, Eq. (9) shows that effective (decoherent) transmission is equal to coherent transmission plus the component of decoherence probes. FIG. S7 shows the *Water* $Na^{+}$ case, and FIG. S8 shows the *Dry* $Na^{+}$ case. Both plots show no significant differences between the trend of decoherent and coherent transmissions. Therefore, we can conclude the same discussions and results as the main manuscript.

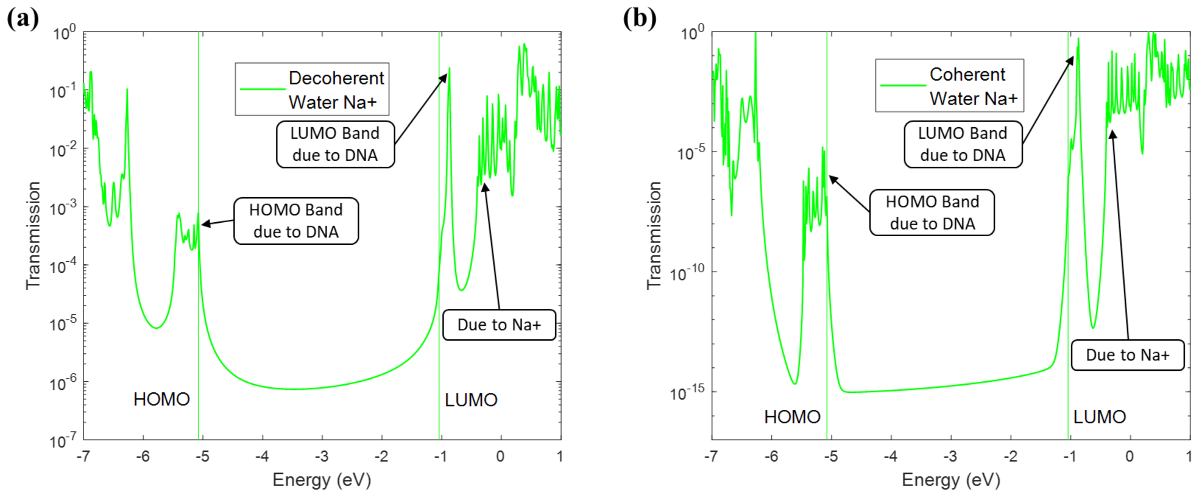


FIG. S7. Transmission of DNA with $Na^{+}$ ions in an implicit water environment. (a) Decoherent transmission, which is the same plot as Fig. 3 (a). (b) Coherent transmission shows the same trend.

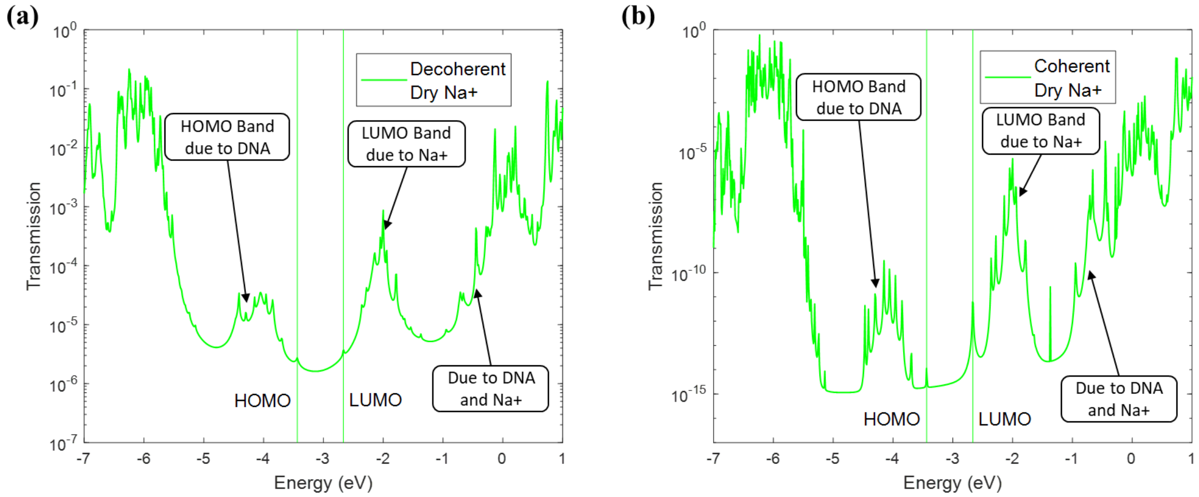


FIG. S8. Transmission of DNA with $Na^{+}$ ions in a dry environment. (a) Decoherent transmission, which is the same plot as Fig. 4 (a). (b) Coherent transmission shows the same trend.

### III. Results for other Sequences

In the main manuscript, our discussion focused on one specific nine-base-pair DNA sequence (5’-CCCGCGCCC-3’). However, our conclusions are not limited to this specific sequence. In order to show that our conclusions can apply to other short-strand DNA, we applied the same simulation procedure as the one discussed in Section II of main manuscript with B3LYP/6-31G(d,p) basis sets for DFT calculation to various DNA strands with different sequences and lengths as shown below:

1. 5’-CCCCCC-3’
   3’-GGGGGG-5’ (Fig. S9 and Fig. S10)
2. 5’-TTTTTT-3’
   3’-AAAAAA-5’ (Fig. S11 and Fig. S12)
3. 5’-CCCTTTCCC-3’
   3’-GGGAAAGGG-5’ (Fig. S13 and Fig. S14)
4. 5’-GGCGCGCGGGCGGGC-3’
   3’-CCGCGCGCCCGCCCG-5’ (Fig. S15 and Fig. S16)
5. 5’-GGCGCAAAAACGGGC-3’
   3’-CCGCGTTTTTGCCCG-5’ (Fig. S17 and Fig. S18)

We chose these sequences based on previously published work. DNA strands 1~3 come from [1]. DNA strands 4~5 come from [2].

The transmission and wavefunction plots are shown in the figure below. For all investigated sequences, the wavefunctions of the HOMO band are localized on Guanine or Adenine bases. These wavefunctions form a transmission channel from left to right using Guanine or Adenine bases. The channel could utilize both strands of a DNA molecule based on the sequence. The HOMO band’s energy levels are consistent with the number of base pairs. With a water solvent, the wavefunctions of the LUMO band are localized on Cytosine or Thymine bases. Under dehydrated condition, the wavefunctions of the LUMO band are localized on $Na^+$ ions. These wavefunctions form two separate channels from left to right (along both backbones of a DNA molecule) using $Na^+$ ions. The LUMO band’s energy levels are consistent with the number of base pairs or $Na^+$ ions, respectively. These results show that regardless of the sequence or length, depending on the dielectric constant of the environment, $Na^+$ ions can significantly impact the charge transport properties of the DNA molecules. Based on the additional calculations, we believe our conclusions can be generalized to other short-strand DNA sequences.

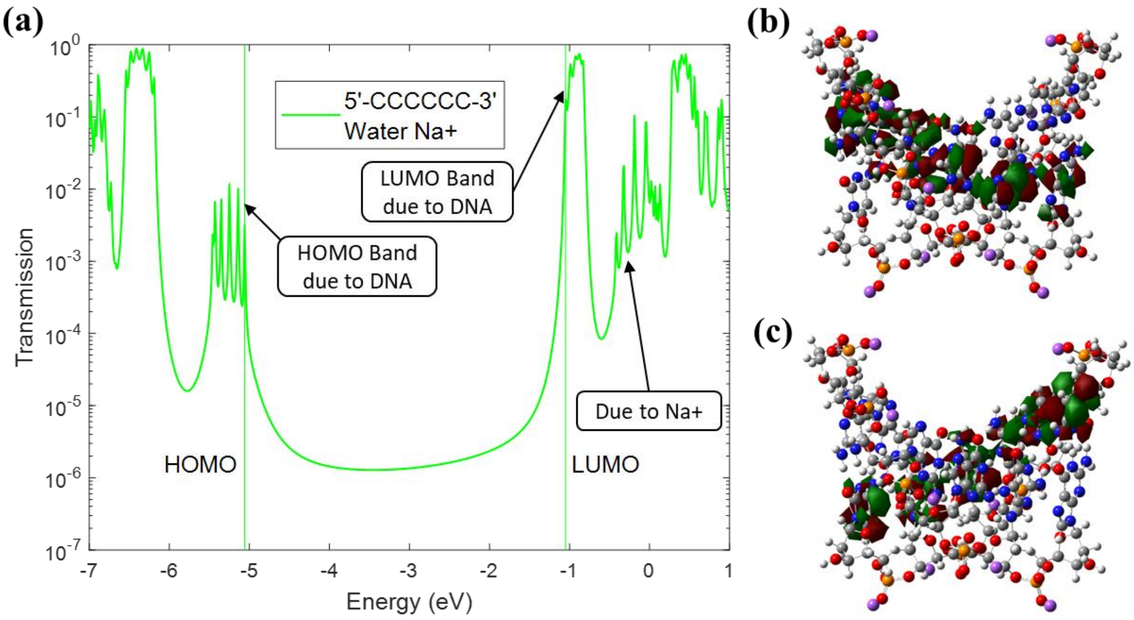


FIG. S9. DFT calculations with 5'-CCCCCC-3' sequence for (a) Decoherent transmission of DNA with $Na^+$ ions in a water environment (*Water* $Na^+$). (b) The wavefunctions of the highest six HOMO energy levels (HOMO band) are localized on Guanine bases. (c) The wavefunctions of the lowest six LUMO energy levels (LUMO band) are localized on Cytosine bases.

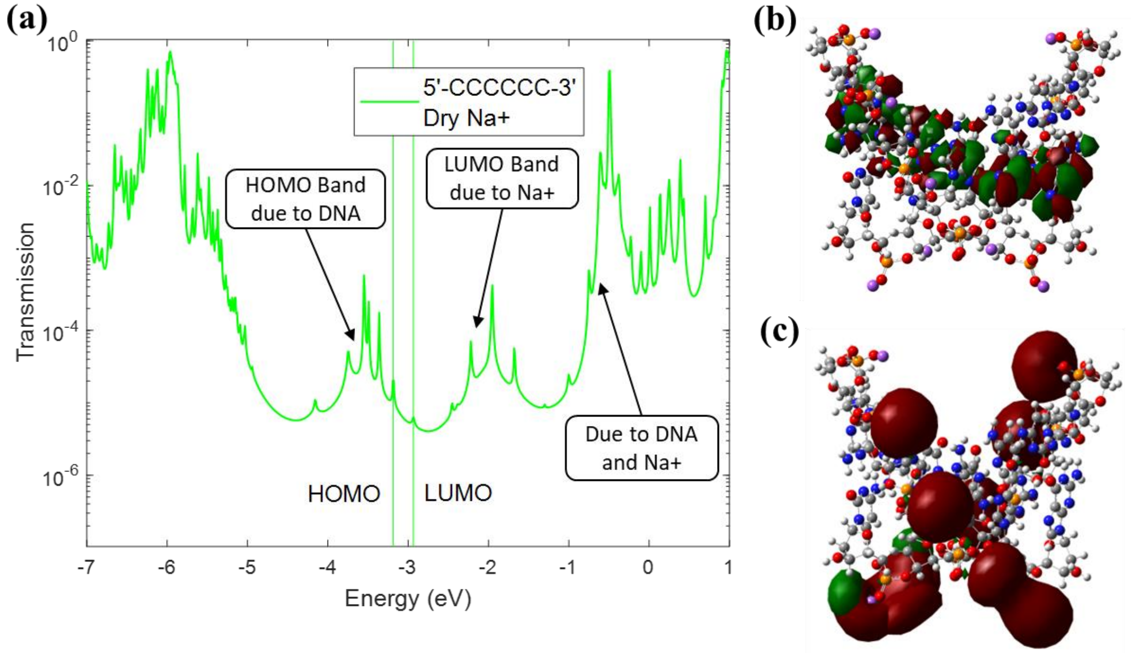


FIG. S10. DFT calculations with 5'-CCCCCC-3' sequence for (a) Decoherent transmission of DNA with $Na^+$ ions in a dry environment (*Dry* $Na^+$). (b) The wavefunctions of the highest six HOMO energy levels (HOMO band) are localized on Guanine bases. (c) The wavefunctions of the lowest ten LUMO energy levels (LUMO band) are localized on $Na^+$ ions.

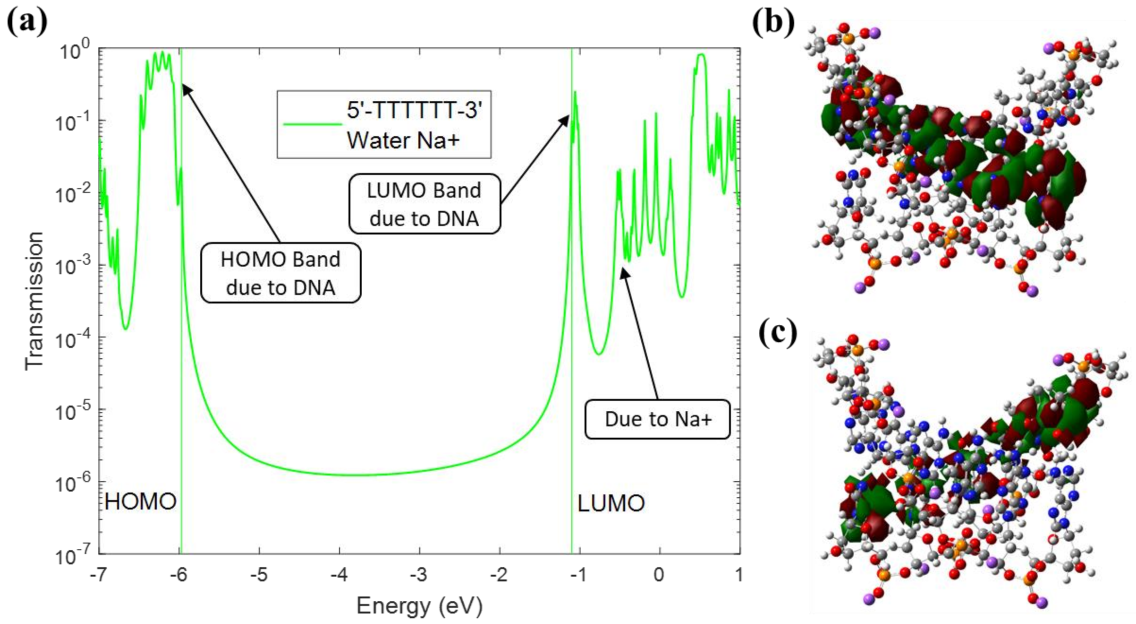


FIG. S11. DFT calculations with 5’-TTTTTT-3’ sequence for (a) Decoherent transmission of DNA with $Na^+$ ions in a water environment (*Water* $Na^+$). (b) The wavefunctions of the highest six HOMO energy levels (HOMO band) are localized on Adenine bases. (c) The wavefunctions of the lowest six LUMO energy levels (LUMO band) are localized on Thymine bases.

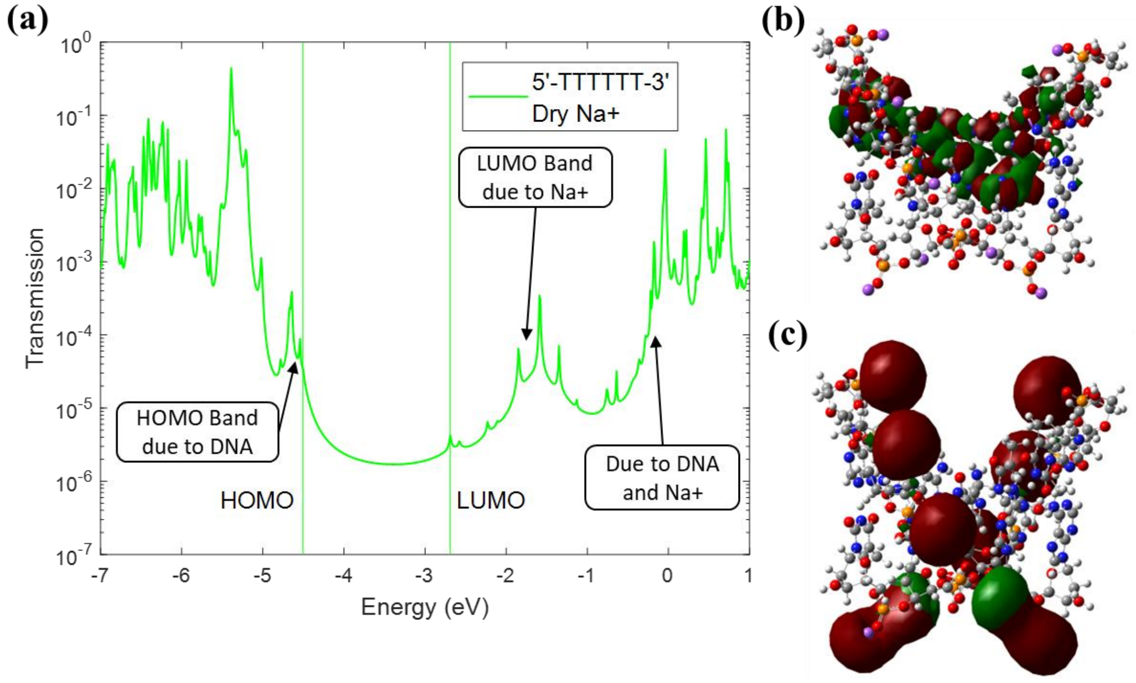


FIG. S12. DFT calculations with 5’-TTTTTT-3’ sequence for (a) Decoherent transmission of DNA with $Na^+$ ions in a dry environment (*Dry* $Na^+$). (b) The wavefunctions of the highest six HOMO energy levels (HOMO band) are localized on Adenine bases. (c) The wavefunctions of the lowest ten LUMO energy levels (LUMO band) are localized on $Na^+$ ions.

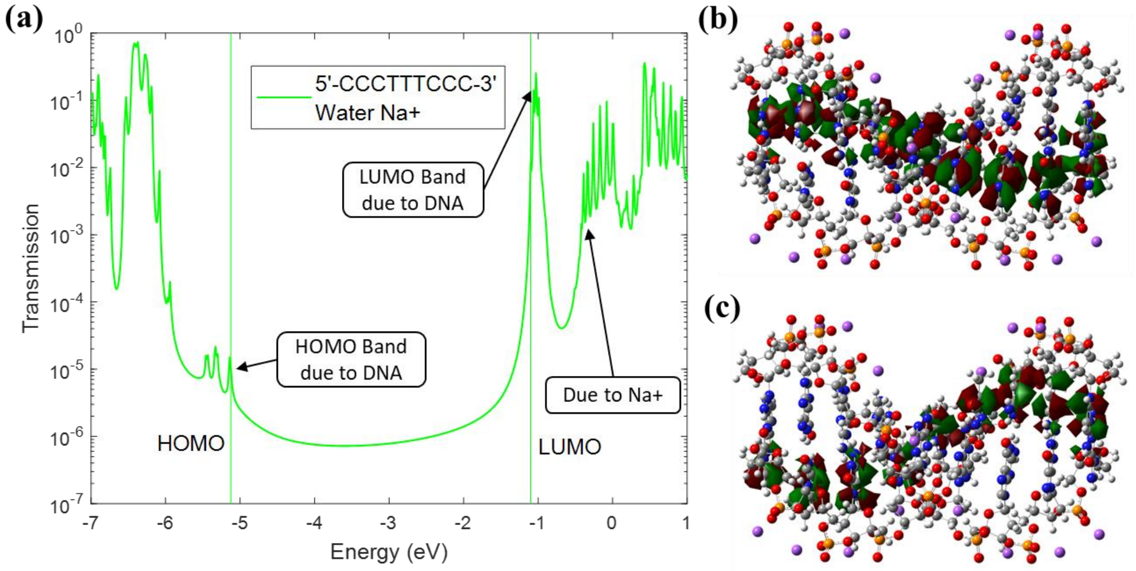


FIG. S13. DFT calculations with 5’-CCCTTTCCC-3’ sequence for (a) Decoherent transmission of DNA with $Na^+$ ions in a water environment (*Water* $Na^+$). (b) The wavefunctions of the highest nine HOMO energy levels (HOMO band) are localized on Guanine or Adenine bases. (c) The wavefunctions of the lowest nine LUMO energy levels (LUMO band) are localized on Cytosine or Thymine bases.

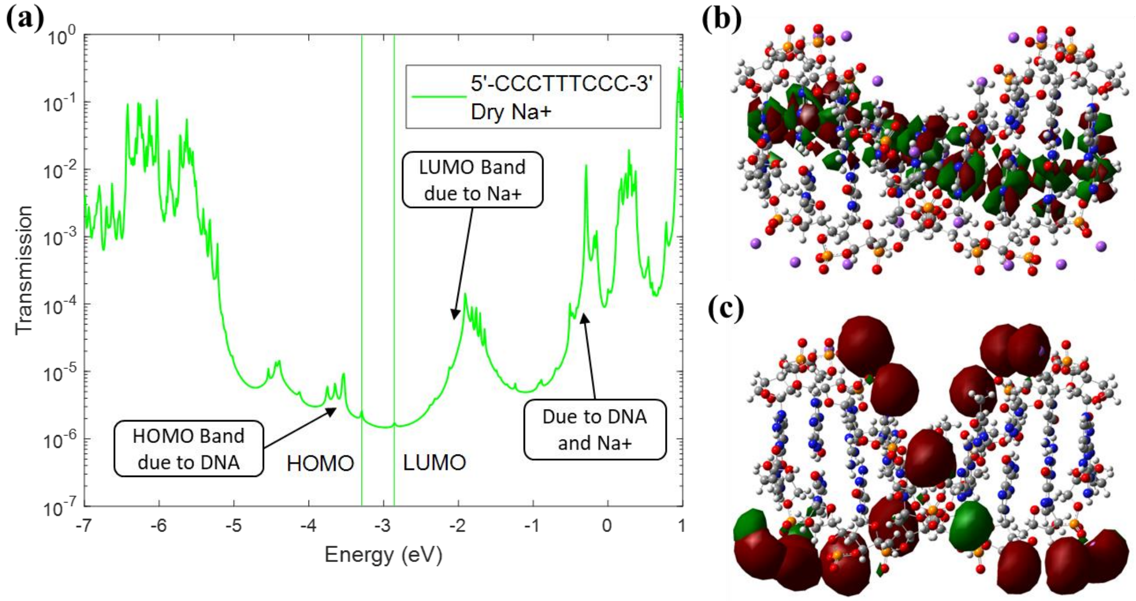


FIG. S14. DFT calculations with 5’-CCCTTTCCC-3’ sequence for (a) Decoherent transmission of DNA with $Na^+$ ions in a dry environment (*Dry* $Na^+$). (b) The wavefunctions of the highest nine HOMO energy levels (HOMO band) are localized on Guanine or Adenine bases. (c) The wavefunctions of the lowest sixteen LUMO energy levels (LUMO band) are localized on $Na^+$ ions.

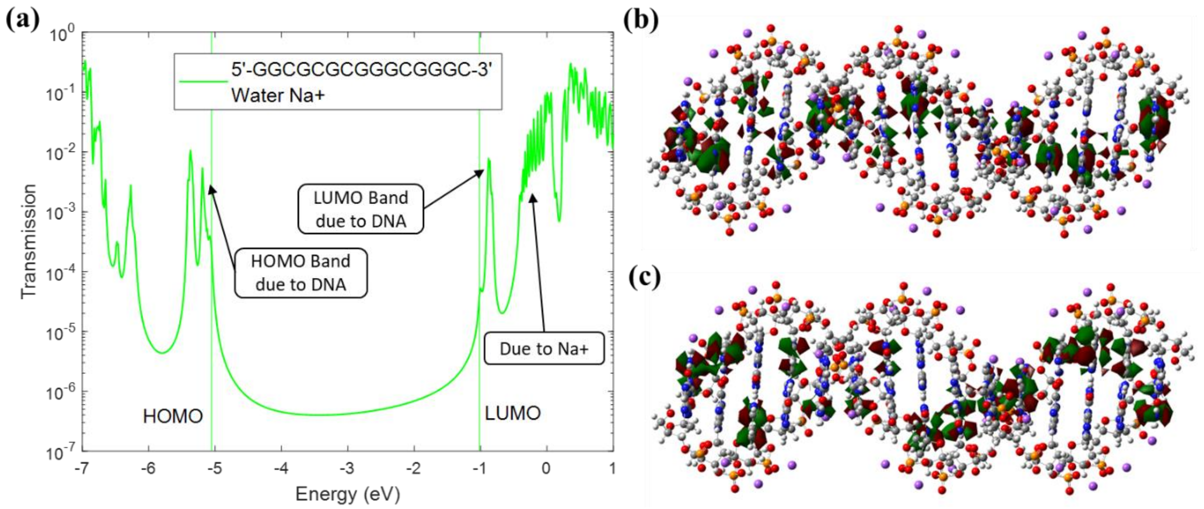


FIG. S15. DFT calculations with 5’-GGCGCGCGGGCGGGC-3’ sequence for (a) Decoherent transmission of DNA with $Na^+$ ions in a water environment (*Water* $Na^+$). (b) The wavefunctions of the highest fifteen HOMO energy levels (HOMO band) are localized on Guanine or Adenine bases. (c) The wavefunctions of the lowest fifteen LUMO energy levels (LUMO band) are localized on Cytosine or Thymine bases.

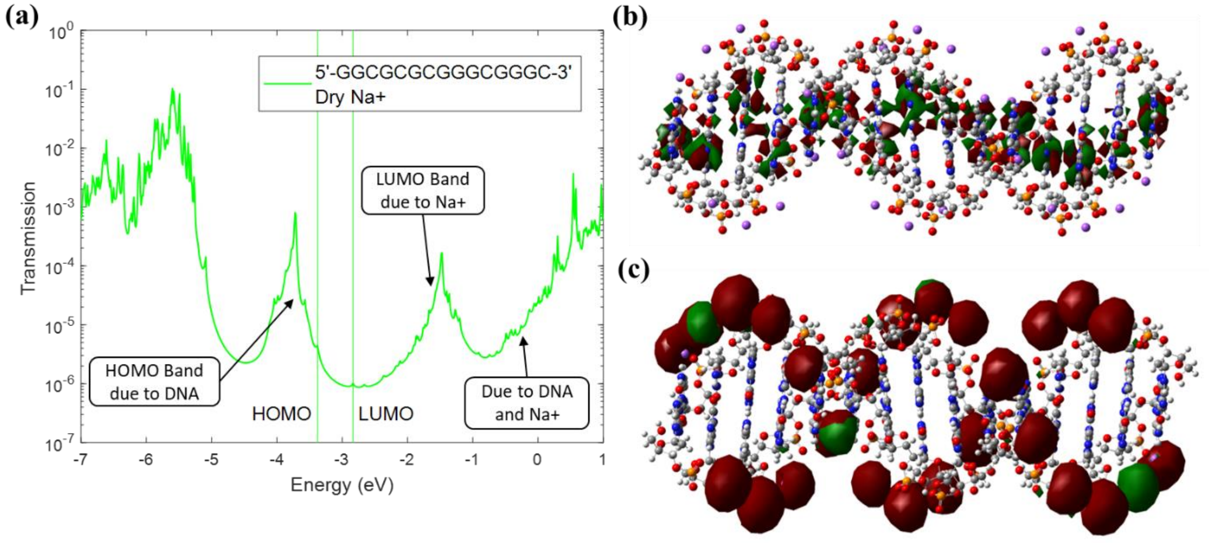


FIG. S16. DFT calculations with 5’-GGCGCGCGGGCGGGC-3’ sequence for (a) Decoherent transmission of DNA with $Na^+$ ions in a dry environment (*Dry* $Na^+$). (b) The wavefunctions of the highest fifteen HOMO energy levels (HOMO band) are localized on Guanine or Adenine bases. (c) The wavefunctions of the lowest twenty-eight LUMO energy levels (LUMO band) are localized on $Na^+$ ions.

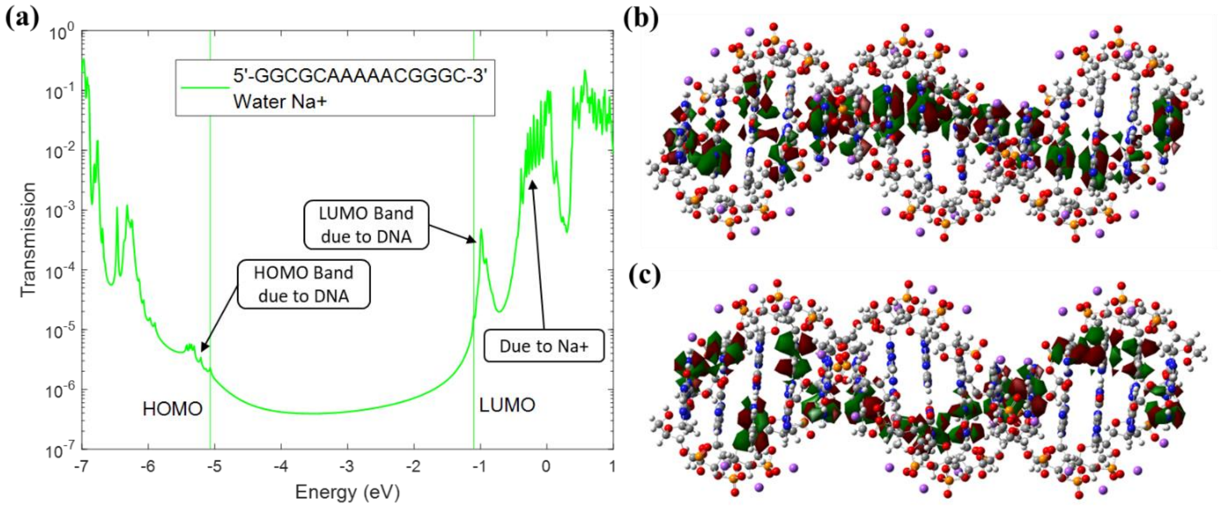


FIG. S17. DFT calculations with 5’-GGCGCAAAAACGGGC-3’ sequence for (a) Decoherent transmission of DNA with $Na^+$ ions in a water environment (*Water* $Na^+$). (b) The wavefunctions of the highest fifteen HOMO energy levels (HOMO band) are localized on Guanine or Adenine bases. (c) The wavefunctions of the lowest fifteen LUMO energy levels (LUMO band) are localized on Cytosine or Thymine bases.

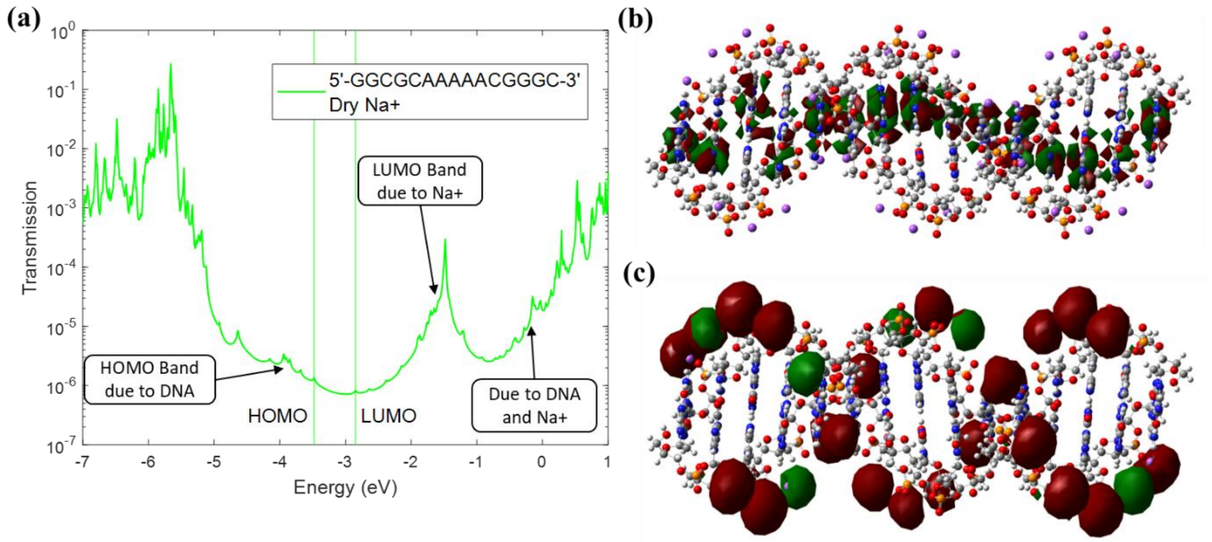


FIG. S18. DFT calculations with 5’-GGCGCAAAAACGGGC-3’ sequence for (a) Decoherent transmission of DNA with $Na^+$ ions in a dry environment (*Dry* $Na^+$). (b) The wavefunctions of the highest fifteen HOMO energy levels (HOMO band) are localized on Guanine or Adenine bases. (c) The wavefunctions of the lowest twenty-eight LUMO energy levels (LUMO band) are localized on $Na^+$ ions.

### IV. Coordinates of the DNA Molecule in Figure 5 of the Main Manuscript

We list each atom's (x, y, z) coordinates for our modeling system below. Sodium ions are located at the end of the list. The sodium ion highlighted in bold is the 7th $Na^+$ ion we removed in the main manuscript (see Fig. 5).

| | | | |
|---|---|---|---|
| H | -0.4230 | -8.1500 | -2.0940 |
| O | 0.4270 | -7.8260 | -1.7880 |
| C | 1.4430 | -7.5100 | -2.7560 |
| H | 1.1000 | -6.7450 | -3.4530 |
| H | 1.6740 | -8.4230 | -3.3050 |
| C | 2.6950 | -7.0200 | -2.0530 |
| H | 3.5740 | -7.0560 | -2.6960 |
| O | 2.4770 | -5.6300 | -1.8230 |
| C | 2.3080 | -5.3330 | -0.4520 |
| H | 3.1120 | -4.6850 | -0.1020 |
| N | 1.0980 | -4.4680 | -0.3700 |
| C | -0.1510 | -5.0050 | -0.3990 |
| H | -0.2760 | -6.0740 | -0.4820 |
| C | -1.2540 | -4.2170 | -0.3250 |
| H | -2.2470 | -4.6650 | -0.3500 |
| C | -1.0350 | -2.8050 | -0.2150 |
| N | -2.0660 | -1.9770 | -0.1380 |
| H | -1.8700 | -0.9890 | -0.0620 |
| H | -3.0100 | -2.3360 | -0.1560 |
| N | 0.1990 | -2.2870 | -0.1870 |
| C | 1.2890 | -3.0930 | -0.2630 |
| O | 2.4450 | -2.6510 | -0.2410 |
| C | 2.9500 | -7.6070 | -0.6650 |
| H | 2.5860 | -8.6290 | -0.5540 |
| C | 2.1480 | -6.6790 | 0.2470 |
| H | 1.3440 | -7.2410 | 0.7220 |
| H | 2.8040 | -6.2670 | 1.0140 |
| O | 4.3240 | -7.5450 | -0.3070 |
| P | 4.7280 | -7.6260 | 1.2390 |
| O | 6.0090 | -8.3560 | 1.3800 |
| O | 3.5920 | -8.1640 | 2.0210 |
| O | 4.9450 | -6.0810 | 1.5920 |
| C | 5.5820 | -5.2270 | 0.6240 |
| H | 4.8550 | -4.8090 | -0.0720 |
| H | 6.3060 | -5.8290 | 0.0750 |
| C | 6.3060 | -4.0950 | 1.3270 |
| H | 7.0380 | -3.6070 | 0.6840 |
| O | 5.3130 | -3.0990 | 1.5570 |
| C | 5.0020 | -2.9580 | 2.9280 |
| H | 5.2720 | -1.9610 | 3.2780 |
| N | 3.5150 | -2.9690 | 3.0100 |
| C | 2.8190 | -4.1380 | 2.9810 |
| H | 3.3460 | -5.0770 | 2.8980 |
| C | 1.4640 | -4.1480 | 3.0550 |
| H | 0.9240 | -5.0950 | 3.0300 |
| C | 0.8110 | -2.8780 | 3.1650 |
| N | -0.5090 | -2.8130 | 3.2420 |
| H | -0.9300 | -1.8980 | 3.3190 |
| H | -1.0620 | -3.6580 | 3.2240 |
| N | 1.5060 | -1.7330 | 3.1930 |
| C | 2.8610 | -1.7440 | 3.1170 |
| O | 3.5360 | -0.7080 | 3.1390 |
| C | 6.8580 | -4.4200 | 2.7150 |
| H | 7.1640 | -5.4600 | 2.8260 |
| C | 5.6640 | -4.1410 | 3.6270 |
| H | 5.3440 | -5.0690 | 4.1020 |
| H | 5.9520 | -3.4220 | 4.3940 |
| O | 7.9330 | -3.5620 | 3.0730 |
| P | 8.3080 | -3.3900 | 4.6190 |
| O | 9.7730 | -3.2280 | 4.7600 |
| O | 7.7050 | -4.4930 | 5.4010 |
| O | 7.5750 | -2.0130 | 4.9720 |
| C | 7.5880 | -0.9480 | 4.0040 |
| H | 6.7540 | -1.0370 | 3.3080 |
| H | 8.5280 | -1.0100 | 3.4550 |
| C | 7.5090 | 0.3940 | 4.7070 |
| H | 7.8150 | 1.2190 | 4.0640 |
| O | 6.1200 | 0.6160 | 4.9370 |
| C | 5.7850 | 0.5470 | 6.3080 |
| H | 5.4170 | 1.5120 | 6.6580 |
| N | 4.5890 | -0.3360 | 6.3900 |
| C | 4.7130 | -1.6900 | 6.3610 |
| H | 5.6910 | -2.1400 | 6.2780 |
| C | 3.6230 | -2.4950 | 6.4350 |
| H | 3.7430 | -3.5780 | 6.4100 |
| C | 2.3480 | -1.8520 | 6.5450 |
| N | 1.2420 | -2.5750 | 6.6220 |
| H | 0.3640 | -2.0830 | 6.6980 |

| | | | |
|---|---|---|---|
| H | 1.2910 | -3.5840 | 6.6040 |
| N | 2.2370 | -0.5170 | 6.5730 |
| C | 3.3400 | 0.2700 | 6.4970 |
| O | 3.2770 | 1.5060 | 6.5190 |
| C | 8.1460 | 0.4550 | 6.0950 |
| H | 9.0050 | -0.2070 | 6.2050 |
| C | 7.0160 | -0.0210 | 7.0070 |
| H | 7.3020 | -0.9590 | 7.4820 |
| H | 6.8260 | 0.7300 | 7.7740 |
| O | 8.5120 | 1.7810 | 6.4530 |
| P | 8.7140 | 2.1400 | 7.9990 |
| O | 9.8040 | 3.1330 | 8.1400 |
| O | 8.8740 | 0.8940 | 8.7810 |
| O | 7.3120 | 2.8240 | 8.3520 |
| C | 6.6960 | 3.6930 | 7.3840 |
| H | 6.0740 | 3.1300 | 6.6890 |
| H | 7.4920 | 4.1950 | 6.8340 |
| C | 5.8430 | 4.7320 | 8.0870 |
| H | 5.6060 | 5.5790 | 7.4430 |
| O | 4.5890 | 4.0960 | 8.3170 |
| C | 4.3590 | 3.8430 | 9.6880 |
| H | 3.4950 | 4.4080 | 10.0390 |
| N | 3.9100 | 2.4250 | 9.7700 |
| C | 4.6630 | 1.2710 | 9.7560 |
| H | 5.7380 | 1.3280 | 9.6750 |
| N | 3.9540 | 0.1710 | 9.8450 |
| C | 2.6400 | 0.6290 | 9.9240 |
| C | 1.4260 | -0.0960 | 10.0370 |
| O | 1.2570 | -1.3110 | 10.0930 |
| N | 0.3190 | 0.7640 | 10.0870 |
| H | -0.5860 | 0.3460 | 10.1680 |
| C | 0.3790 | 2.1430 | 10.0340 |
| N | -0.7910 | 2.7830 | 10.0970 |
| H | -1.7090 | 2.3670 | 10.1670 |
| H | -0.7570 | 3.7910 | 10.0500 |
| N | 1.5190 | 2.8220 | 9.9280 |
| C | 2.6030 | 2.0030 | 9.8790 |
| C | 6.3230 | 5.1570 | 9.4750 |
| H | 7.4070 | 5.1270 | 9.5860 |
| C | 5.6880 | 4.1070 | 10.3870 |
| H | 6.4710 | 3.5160 | 10.8620 |
| H | 5.0930 | 4.6030 | 11.1540 |
| O | 5.8390 | 6.4440 | 9.8330 |
| P | 5.7920 | 6.8540 | 11.3790 |
| O | 6.0900 | 8.2970 | 11.5200 |
| O | 6.6540 | 5.9390 | 12.1610 |
| O | 4.2550 | 6.5820 | 11.7320 |
| C | 3.2460 | 6.9240 | 10.7640 |
| H | 3.0730 | 6.1030 | 10.0680 |
| H | 3.5950 | 7.7980 | 10.2150 |
| C | 1.9460 | 7.2630 | 11.4670 |
| H | 1.2560 | 7.8090 | 10.8230 |
| O | 1.3050 | 6.0110 | 11.6970 |
| C | 1.2680 | 5.6710 | 13.0680 |
| H | 0.2370 | 5.6200 | 13.4180 |
| N | 1.7380 | 4.2600 | 13.1500 |
| C | 3.0640 | 3.9600 | 13.1210 |
| H | 3.7940 | 4.7520 | 13.0380 |
| C | 3.4930 | 2.6740 | 13.1950 |
| H | 4.5600 | 2.4520 | 13.1700 |
| C | 2.4860 | 1.6610 | 13.3050 |
| N | 2.8330 | 0.3850 | 13.3820 |
| H | 2.0940 | -0.2990 | 13.4590 |
| H | 3.8080 | 0.1200 | 13.3640 |
| N | 1.1830 | 1.9680 | 13.3330 |
| C | 0.7750 | 3.2600 | 13.2570 |
| O | -0.4190 | 3.5820 | 13.2790 |
| C | 2.0840 | 7.8880 | 12.8550 |
| H | 2.9780 | 8.5010 | 12.9660 |
| C | 2.1880 | 6.6660 | 13.7670 |
| H | 3.1690 | 6.6490 | 14.2420 |
| H | 1.4150 | 6.7170 | 14.5330 |
| O | 0.9360 | 8.6450 | 13.2130 |
| P | 0.6570 | 8.9490 | 14.7590 |
| O | 0.0500 | 10.2920 | 14.9000 |
| O | 1.8920 | 8.7160 | 15.5410 |
| O | -0.4270 | 7.8260 | 15.1120 |
| C | -1.4430 | 7.5100 | 14.1440 |
| H | -1.1000 | 6.7450 | 13.4470 |
| H | -1.6740 | 8.4230 | 13.5950 |
| C | -2.6950 | 7.0200 | 14.8470 |
| H | -3.5740 | 7.0560 | 14.2040 |
| O | -2.4770 | 5.6300 | 15.0770 |
| C | -2.3080 | 5.3330 | 16.4480 |
| H | -3.1120 | 4.6860 | 16.7990 |
| N | -1.0980 | 4.4680 | 16.5300 |
| C | 0.2320 | 4.8270 | 16.5160 |
| H | 0.5090 | 5.8680 | 16.4350 |

| | | | |
|---|---|---|---|
| N | 1.0590 | 3.8140 | 16.6050 |
| C | 0.2180 | 2.7050 | 16.6840 |
| C | 0.5320 | 1.3260 | 16.7970 |
| O | 1.6350 | 0.7900 | 16.8530 |
| N | -0.6280 | 0.5400 | 16.8470 |
| H | -0.5090 | -0.4500 | 16.9290 |
| C | -1.9210 | 1.0230 | 16.7940 |
| N | -2.8910 | 0.1080 | 16.8570 |
| H | -2.7780 | -0.8930 | 16.9270 |
| H | -3.8390 | 0.4520 | 16.8100 |
| N | -2.2140 | 2.3170 | 16.6880 |
| C | -1.1010 | 3.0940 | 16.6390 |
| C | -2.9500 | 7.6070 | 16.2350 |
| H | -2.5860 | 8.6290 | 16.3460 |
| C | -2.1480 | 6.6790 | 17.1470 |
| H | -1.3440 | 7.2410 | 17.6220 |
| H | -2.8040 | 6.2670 | 17.9140 |
| O | -4.3240 | 7.5450 | 16.5930 |
| P | -4.7280 | 7.6260 | 18.1390 |
| O | -6.0090 | 8.3560 | 18.2800 |
| O | -3.5920 | 8.1640 | 18.9210 |
| O | -4.9450 | 6.0810 | 18.4920 |
| C | -5.5820 | 5.2270 | 17.5240 |
| H | -4.8550 | 4.8090 | 16.8280 |
| H | -6.3060 | 5.8290 | 16.9750 |
| C | -6.3060 | 4.0950 | 18.2270 |
| H | -7.0380 | 3.6070 | 17.5840 |
| O | -5.3130 | 3.0990 | 18.4570 |
| C | -5.0020 | 2.9580 | 19.8280 |
| H | -5.2720 | 1.9610 | 20.1780 |
| N | -3.5150 | 2.9690 | 19.9100 |
| C | -2.8190 | 4.1380 | 19.8810 |
| H | -3.3460 | 5.0770 | 19.7980 |
| C | -1.4640 | 4.1480 | 19.9550 |
| H | -0.9240 | 5.0950 | 19.9300 |
| C | -0.8110 | 2.8780 | 20.0650 |
| N | 0.5090 | 2.8130 | 20.1420 |
| H | 0.9300 | 1.8980 | 20.2190 |
| H | 1.0620 | 3.6580 | 20.1240 |
| N | -1.5060 | 1.7330 | 20.0930 |
| C | -2.8610 | 1.7440 | 20.0170 |
| O | -3.5360 | 0.7080 | 20.0390 |
| C | -6.8580 | 4.4200 | 19.6150 |
| H | -7.1640 | 5.4600 | 19.7260 |

| | | | |
|---|---|---|---|
| C | -5.6640 | 4.1410 | 20.5270 |
| H | -5.3440 | 5.0690 | 21.0020 |
| H | -5.9520 | 3.4220 | 21.2940 |
| O | -7.9330 | 3.5620 | 19.9730 |
| P | -8.3080 | 3.3900 | 21.5190 |
| O | -9.7730 | 3.2280 | 21.6600 |
| O | -7.7050 | 4.4930 | 22.3010 |
| O | -7.5750 | 2.0130 | 21.8720 |
| C | -7.5880 | 0.9480 | 20.9040 |
| H | -6.7540 | 1.0370 | 20.2080 |
| H | -8.5280 | 1.0100 | 20.3550 |
| C | -7.5090 | -0.3940 | 21.6070 |
| H | -7.8150 | -1.2190 | 20.9640 |
| O | -6.1200 | -0.6160 | 21.8370 |
| C | -5.7850 | -0.5470 | 23.2080 |
| H | -5.4170 | -1.5120 | 23.5580 |
| N | -4.5890 | 0.3360 | 23.2900 |
| C | -4.7130 | 1.6900 | 23.2610 |
| H | -5.6910 | 2.1400 | 23.1780 |
| C | -3.6230 | 2.4950 | 23.3350 |
| H | -3.7430 | 3.5780 | 23.3100 |
| C | -2.3480 | 1.8520 | 23.4450 |
| N | -1.2420 | 2.5750 | 23.5220 |
| H | -0.3640 | 2.0830 | 23.5980 |
| H | -1.2910 | 3.5840 | 23.5040 |
| N | -2.2370 | 0.5170 | 23.4730 |
| C | -3.3400 | -0.2700 | 23.3970 |
| O | -3.2770 | -1.5060 | 23.4190 |
| C | -8.1460 | -0.4550 | 22.9950 |
| H | -9.0050 | 0.2070 | 23.1050 |
| C | -7.0160 | 0.0210 | 23.9070 |
| H | -7.3020 | 0.9590 | 24.3820 |
| H | -6.8260 | -0.7300 | 24.6740 |
| O | -8.5120 | -1.7810 | 23.3530 |
| P | -8.7140 | -2.1400 | 24.8990 |
| O | -9.8040 | -3.1330 | 25.0400 |
| O | -8.8740 | -0.8940 | 25.6810 |
| O | -7.3120 | -2.8240 | 25.2520 |
| C | -6.6960 | -3.6930 | 24.2840 |
| H | -6.0740 | -3.1300 | 23.5890 |
| H | -7.4920 | -4.1950 | 23.7340 |
| C | -5.8430 | -4.7320 | 24.9870 |
| H | -5.6060 | -5.5790 | 24.3430 |
| O | -4.5890 | -4.0960 | 25.2170 |

| | | | |
|---|---|---|---|
| C | -4.3590 | -3.8430 | 26.5880 |
| H | -3.4950 | -4.4070 | 26.9380 |
| N | -3.9100 | -2.4250 | 26.6700 |
| C | -4.8060 | -1.4030 | 26.6410 |
| H | -5.8620 | -1.6150 | 26.5580 |
| C | -4.3980 | -0.1110 | 26.7150 |
| H | -5.1320 | 0.6950 | 26.6900 |
| C | -2.9880 | 0.1180 | 26.8250 |
| N | -2.5180 | 1.3540 | 26.9020 |
| H | -1.5180 | 1.4720 | 26.9780 |
| H | -3.1500 | 2.1410 | 26.8840 |
| N | -2.1140 | -0.8960 | 26.8530 |
| C | -2.5430 | -2.1820 | 26.7770 |
| O | -1.7660 | -3.1440 | 26.7990 |
| C | -6.3230 | -5.1570 | 26.3750 |
| H | -7.4070 | -5.1270 | 26.4860 |
| C | -5.6880 | -4.1070 | 27.2870 |
| H | -6.4710 | -3.5160 | 27.7620 |
| H | -5.0930 | -4.6030 | 28.0540 |
| O | -5.8390 | -6.4440 | 26.7330 |
| H | -6.1150 | -6.7540 | 27.5990 |
| H | 7.6200 | 2.9210 | 29.1350 |
| O | 7.5750 | 2.0130 | 28.8280 |
| C | 7.5880 | 0.9480 | 29.7960 |
| H | 6.7540 | 1.0370 | 30.4920 |
| H | 8.5280 | 1.0100 | 30.3450 |
| C | 7.5090 | -0.3940 | 29.0930 |
| H | 7.8150 | -1.2190 | 29.7360 |
| O | 6.1200 | -0.6160 | 28.8630 |
| C | 5.7850 | -0.5470 | 27.4920 |
| H | 5.4180 | -1.5120 | 27.1410 |
| N | 4.5890 | 0.3360 | 27.4100 |
| C | 4.5200 | 1.7120 | 27.4240 |
| H | 5.4240 | 2.2970 | 27.5050 |
| N | 3.3000 | 2.1860 | 27.3350 |
| C | 2.5060 | 1.0430 | 27.2560 |
| C | 1.0970 | 0.9160 | 27.1430 |
| O | 0.2460 | 1.7990 | 27.0870 |
| N | 0.7070 | -0.4300 | 27.0930 |
| H | -0.2710 | -0.6230 | 27.0110 |
| C | 1.5660 | -1.5100 | 27.1460 |
| N | 0.9960 | -2.7160 | 27.0830 |
| H | 0.0090 | -2.9190 | 27.0130 |
| H | 1.6170 | -3.5110 | 27.1300 |
| N | 2.8880 | -1.3900 | 27.2520 |
| C | 3.2830 | -0.0910 | 27.3010 |
| C | 8.1460 | -0.4550 | 27.7050 |
| H | 9.0050 | 0.2070 | 27.5950 |
| C | 7.0160 | 0.0210 | 26.7930 |
| H | 7.3020 | 0.9590 | 26.3180 |
| H | 6.8260 | -0.7300 | 26.0260 |
| O | 8.5120 | -1.7810 | 27.3470 |
| P | 8.7140 | -2.1400 | 25.8010 |
| O | 9.8040 | -3.1330 | 25.6600 |
| O | 8.8740 | -0.8940 | 25.0190 |
| O | 7.3120 | -2.8240 | 25.4480 |
| C | 6.6960 | -3.6930 | 26.4160 |
| H | 6.0740 | -3.1300 | 27.1110 |
| H | 7.4920 | -4.1950 | 26.9660 |
| C | 5.8430 | -4.7320 | 25.7130 |
| H | 5.6060 | -5.5790 | 26.3570 |
| O | 4.5890 | -4.0960 | 25.4830 |
| C | 4.3590 | -3.8430 | 24.1120 |
| H | 3.4950 | -4.4080 | 23.7610 |
| N | 3.9100 | -2.4250 | 24.0300 |
| C | 4.6630 | -1.2710 | 24.0440 |
| H | 5.7380 | -1.3280 | 24.1250 |
| N | 3.9540 | -0.1710 | 23.9550 |
| C | 2.6400 | -0.6290 | 23.8760 |
| C | 1.4260 | 0.0960 | 23.7630 |
| O | 1.2570 | 1.3110 | 23.7070 |
| N | 0.3190 | -0.7640 | 23.7130 |
| H | -0.5860 | -0.3460 | 23.6320 |
| C | 0.3790 | -2.1430 | 23.7660 |
| N | -0.7910 | -2.7830 | 23.7030 |
| H | -1.7090 | -2.3670 | 23.6330 |
| H | -0.7570 | -3.7910 | 23.7500 |
| N | 1.5190 | -2.8220 | 23.8720 |
| C | 2.6030 | -2.0030 | 23.9210 |
| C | 6.3230 | -5.1570 | 24.3250 |
| H | 7.4070 | -5.1270 | 24.2140 |
| C | 5.6880 | -4.1070 | 23.4130 |
| H | 6.4710 | -3.5160 | 22.9380 |
| H | 5.0930 | -4.6030 | 22.6460 |
| O | 5.8390 | -6.4440 | 23.9670 |
| P | 5.7920 | -6.8540 | 22.4210 |
| O | 6.0900 | -8.2970 | 22.2800 |
| O | 6.6540 | -5.9390 | 21.6390 |

| | | | |
|---|---|---|---|
| O | 4.2550 | -6.5820 | 22.0680 |
| C | 3.2460 | -6.9240 | 23.0360 |
| H | 3.0730 | -6.1030 | 23.7320 |
| H | 3.5950 | -7.7980 | 23.5850 |
| C | 1.9460 | -7.2630 | 22.3330 |
| H | 1.2560 | -7.8090 | 22.9770 |
| O | 1.3050 | -6.0110 | 22.1030 |
| C | 1.2680 | -5.6710 | 20.7320 |
| H | 0.2370 | -5.6200 | 20.3800 |
| N | 1.7380 | -4.2600 | 20.6500 |
| C | 3.0250 | -3.7690 | 20.6640 |
| H | 3.8610 | -4.4480 | 20.7450 |
| N | 3.0980 | -2.4630 | 20.5750 |
| C | 1.7660 | -2.0610 | 20.4960 |
| C | 1.2100 | -0.7600 | 20.3830 |
| O | 1.7870 | 0.3220 | 20.3270 |
| N | -0.1910 | -0.8060 | 20.3330 |
| H | -0.6770 | 0.0640 | 20.2510 |
| C | -0.9530 | -1.9560 | 20.3860 |
| N | -2.2750 | -1.7870 | 20.3230 |
| H | -2.7730 | -0.9110 | 20.2530 |
| H | -2.8390 | -2.6230 | 20.3700 |
| N | -0.4290 | -3.1760 | 20.4920 |
| C | 0.9280 | -3.1500 | 20.5410 |
| C | 2.0840 | -7.8880 | 20.9450 |
| H | 2.9780 | -8.5010 | 20.8340 |
| C | 2.1880 | -6.6660 | 20.0330 |
| H | 3.1690 | -6.6490 | 19.5580 |
| H | 1.4150 | -6.7170 | 19.2670 |
| O | 0.9360 | -8.6450 | 20.5870 |
| P | 0.6570 | -8.9490 | 19.0410 |
| O | 0.0500 | -10.2920 | 18.9000 |
| O | 1.8920 | -8.7160 | 18.2590 |
| O | -0.4270 | -7.8260 | 18.6880 |
| C | -1.4430 | -7.5100 | 19.6560 |
| H | -1.1000 | -6.7450 | 20.3530 |
| H | -1.6740 | -8.4230 | 20.2050 |
| C | -2.6950 | -7.0200 | 18.9530 |
| H | -3.5740 | -7.0560 | 19.5960 |
| O | -2.4770 | -5.6300 | 18.7230 |
| C | -2.3080 | -5.3330 | 17.3520 |
| H | -3.1120 | -4.6850 | 17.0020 |
| N | -1.0980 | -4.4680 | 17.2700 |
| C | 0.1510 | -5.0050 | 17.2990 |

| | | | |
|---|---|---|---|
| H | 0.2760 | -6.0740 | 17.3820 |
| C | 1.2540 | -4.2170 | 17.2250 |
| H | 2.2470 | -4.6650 | 17.2500 |
| C | 1.0350 | -2.8050 | 17.1150 |
| N | 2.0660 | -1.9770 | 17.0380 |
| H | 1.8700 | -0.9890 | 16.9620 |
| H | 3.0100 | -2.3360 | 17.0560 |
| N | -0.1990 | -2.2870 | 17.0870 |
| C | -1.2890 | -3.0930 | 17.1630 |
| O | -2.4450 | -2.6510 | 17.1410 |
| C | -2.9500 | -7.6070 | 17.5650 |
| H | -2.5860 | -8.6290 | 17.4540 |
| C | -2.1480 | -6.6790 | 16.6530 |
| H | -1.3440 | -7.2410 | 16.1780 |
| H | -2.8040 | -6.2670 | 15.8860 |
| O | -4.3240 | -7.5450 | 17.2070 |
| P | -4.7280 | -7.6260 | 15.6610 |
| O | -6.0090 | -8.3560 | 15.5200 |
| O | -3.5920 | -8.1640 | 14.8790 |
| O | -4.9450 | -6.0810 | 15.3080 |
| C | -5.5820 | -5.2270 | 16.2760 |
| H | -4.8550 | -4.8090 | 16.9720 |
| H | -6.3060 | -5.8290 | 16.8250 |
| C | -6.3060 | -4.0950 | 15.5730 |
| H | -7.0380 | -3.6070 | 16.2160 |
| O | -5.3130 | -3.0990 | 15.3430 |
| C | -5.0020 | -2.9580 | 13.9720 |
| H | -5.2720 | -1.9620 | 13.6210 |
| N | -3.5150 | -2.9690 | 13.8900 |
| C | -2.6500 | -4.0420 | 13.9040 |
| H | -3.0370 | -5.0470 | 13.9850 |
| N | -1.3850 | -3.7080 | 13.8150 |
| C | -1.4140 | -2.3170 | 13.7360 |
| C | -0.3490 | -1.3860 | 13.6230 |
| O | 0.8580 | -1.6000 | 13.5670 |
| N | -0.8250 | -0.0680 | 13.5730 |
| H | -0.1480 | 0.6630 | 13.4920 |
| C | -2.1550 | 0.3010 | 13.6260 |
| N | -2.4020 | 1.6120 | 13.5630 |
| H | -1.7220 | 2.3560 | 13.4930 |
| H | -3.3720 | 1.8910 | 13.6100 |
| N | -3.1530 | -0.5730 | 13.7320 |
| C | -2.7090 | -1.8560 | 13.7810 |
| C | -6.8580 | -4.4200 | 14.1850 |

| | | | |
|---|---|---|---|
| H | -7.1640 | -5.4600 | 14.0740 |
| C | -5.6640 | -4.1410 | 13.2730 |
| H | -5.3440 | -5.0690 | 12.7980 |
| H | -5.9520 | -3.4220 | 12.5060 |
| O | -7.9330 | -3.5620 | 13.8270 |
| P | -8.3080 | -3.3900 | 12.2810 |
| O | -9.7730 | -3.2280 | 12.1400 |
| O | -7.7050 | -4.4930 | 11.4990 |
| O | -7.5750 | -2.0130 | 11.9280 |
| C | -7.5880 | -0.9480 | 12.8960 |
| H | -6.7540 | -1.0370 | 13.5920 |
| H | -8.5280 | -1.0100 | 13.4450 |
| C | -7.5090 | 0.3940 | 12.1930 |
| H | -7.8150 | 1.2190 | 12.8360 |
| O | -6.1200 | 0.6160 | 11.9630 |
| C | -5.7850 | 0.5470 | 10.5920 |
| H | -5.4170 | 1.5120 | 10.2420 |
| N | -4.5890 | -0.3360 | 10.5100 |
| C | -4.7130 | -1.6900 | 10.5390 |
| H | -5.6910 | -2.1400 | 10.6220 |
| C | -3.6230 | -2.4950 | 10.4650 |
| H | -3.7430 | -3.5780 | 10.4900 |
| C | -2.3480 | -1.8520 | 10.3550 |
| N | -1.2420 | -2.5750 | 10.2780 |
| H | -0.3640 | -2.0830 | 10.2020 |
| H | -1.2910 | -3.5840 | 10.2960 |
| N | -2.2370 | -0.5170 | 10.3270 |
| C | -3.3400 | 0.2700 | 10.4030 |
| O | -3.2770 | 1.5060 | 10.3810 |
| C | -8.1460 | 0.4550 | 10.8050 |
| H | -9.0050 | -0.2070 | 10.6950 |
| C | -7.0160 | -0.0210 | 9.8930 |
| H | -7.3020 | -0.9590 | 9.4180 |
| H | -6.8260 | 0.7300 | 9.1260 |
| O | -8.5120 | 1.7810 | 10.4470 |
| P | -8.7140 | 2.1400 | 8.9010 |
| O | -9.8040 | 3.1330 | 8.7600 |
| O | -8.8740 | 0.8940 | 8.1190 |
| O | -7.3120 | 2.8240 | 8.5480 |
| C | -6.6960 | 3.6930 | 9.5160 |
| H | -6.0740 | 3.1300 | 10.2110 |
| H | -7.4920 | 4.1950 | 10.0660 |
| C | -5.8430 | 4.7320 | 8.8130 |
| H | -5.6060 | 5.5790 | 9.4570 |

| | | | |
|---|---|---|---|
| O | -4.5890 | 4.0960 | 8.5830 |
| C | -4.3590 | 3.8430 | 7.2120 |
| H | -3.4950 | 4.4080 | 6.8610 |
| N | -3.9100 | 2.4250 | 7.1300 |
| C | -4.6630 | 1.2710 | 7.1440 |
| H | -5.7380 | 1.3280 | 7.2250 |
| N | -3.9540 | 0.1710 | 7.0550 |
| C | -2.6400 | 0.6290 | 6.9760 |
| C | -1.4260 | -0.0960 | 6.8630 |
| O | -1.2570 | -1.3110 | 6.8070 |
| N | -0.3190 | 0.7640 | 6.8130 |
| H | 0.5860 | 0.3460 | 6.7320 |
| C | -0.3790 | 2.1430 | 6.8660 |
| N | 0.7910 | 2.7830 | 6.8030 |
| H | 1.7090 | 2.3670 | 6.7330 |
| H | 0.7570 | 3.7910 | 6.8500 |
| N | -1.5190 | 2.8220 | 6.9720 |
| C | -2.6030 | 2.0030 | 7.0210 |
| C | -6.3230 | 5.1570 | 7.4250 |
| H | -7.4070 | 5.1270 | 7.3140 |
| C | -5.6880 | 4.1070 | 6.5130 |
| H | -6.4710 | 3.5160 | 6.0380 |
| H | -5.0930 | 4.6030 | 5.7460 |
| O | -5.8390 | 6.4440 | 7.0670 |
| P | -5.7920 | 6.8540 | 5.5210 |
| O | -6.0900 | 8.2970 | 5.3800 |
| O | -6.6540 | 5.9390 | 4.7390 |
| O | -4.2550 | 6.5820 | 5.1680 |
| C | -3.2460 | 6.9240 | 6.1360 |
| H | -3.0730 | 6.1030 | 6.8320 |
| H | -3.5950 | 7.7980 | 6.6850 |
| C | -1.9460 | 7.2630 | 5.4330 |
| H | -1.2560 | 7.8090 | 6.0770 |
| O | -1.3050 | 6.0110 | 5.2030 |
| C | -1.2680 | 5.6710 | 3.8320 |
| H | -0.2370 | 5.6200 | 3.4800 |
| N | -1.7380 | 4.2600 | 3.7500 |
| C | -3.0250 | 3.7690 | 3.7640 |
| H | -3.8610 | 4.4480 | 3.8450 |
| N | -3.0980 | 2.4630 | 3.6750 |
| C | -1.7660 | 2.0610 | 3.5960 |
| C | -1.2100 | 0.7600 | 3.4830 |
| O | -1.7870 | -0.3220 | 3.4270 |
| N | 0.1910 | 0.8060 | 3.4330 |

| | | | |
|---|---|---|---|
| H | 0.6770 | -0.0640 | 3.3510 |
| C | 0.9530 | 1.9560 | 3.4860 |
| N | 2.2750 | 1.7870 | 3.4230 |
| H | 2.7730 | 0.9110 | 3.3530 |
| H | 2.8390 | 2.6230 | 3.4700 |
| N | 0.4290 | 3.1760 | 3.5920 |
| C | -0.9280 | 3.1500 | 3.6410 |
| C | -2.0840 | 7.8880 | 4.0450 |
| H | -2.9780 | 8.5010 | 3.9340 |
| C | -2.1880 | 6.6660 | 3.1330 |
| H | -3.1690 | 6.6490 | 2.6580 |
| H | -1.4150 | 6.7170 | 2.3670 |
| O | -0.9360 | 8.6450 | 3.6870 |
| P | -0.6570 | 8.9490 | 2.1410 |
| O | -0.0500 | 10.2920 | 2.0000 |
| O | -1.8920 | 8.7160 | 1.3590 |
| O | 0.4270 | 7.8260 | 1.7880 |
| C | 1.4430 | 7.5100 | 2.7560 |
| H | 1.1000 | 6.7450 | 3.4530 |
| H | 1.6740 | 8.4230 | 3.3050 |
| C | 2.6950 | 7.0200 | 2.0530 |
| H | 3.5740 | 7.0560 | 2.6960 |
| O | 2.4770 | 5.6300 | 1.8230 |
| C | 2.3080 | 5.3330 | 0.4520 |
| H | 3.1120 | 4.6860 | 0.1010 |
| N | 1.0980 | 4.4680 | 0.3700 |
| C | -0.2320 | 4.8270 | 0.3840 |
| H | -0.5090 | 5.8680 | 0.4650 |
| N | -1.0590 | 3.8140 | 0.2950 |
| C | -0.2180 | 2.7050 | 0.2160 |
| C | -0.5320 | 1.3260 | 0.1030 |
| O | -1.6350 | 0.7900 | 0.0470 |
| N | 0.6280 | 0.5400 | 0.0530 |
| H | 0.5090 | -0.4500 | -0.0290 |
| C | 1.9210 | 1.0230 | 0.1060 |
| N | 2.8910 | 0.1080 | 0.0430 |
| H | 2.7780 | -0.8930 | -0.0270 |
| H | 3.8390 | 0.4520 | 0.0900 |
| N | 2.2140 | 2.3170 | 0.2120 |
| C | 1.1010 | 3.0940 | 0.2610 |
| C | 2.9500 | 7.6070 | 0.6650 |
| H | 2.5860 | 8.6290 | 0.5540 |
| C | 2.1480 | 6.6790 | -0.2470 |
| H | 1.3440 | 7.2410 | -0.7220 |
| H | 2.8040 | 6.2670 | -1.0140 |
| O | 4.3240 | 7.5450 | 0.3070 |
| H | 4.5340 | 7.9030 | -0.5590 |
| Na | 5.2048 | -8.0672 | 3.8233 |
| Na | 8.5147 | -3.3530 | 7.3796 |
| Na | 8.8594 | 2.2923 | 10.7596 |
| Na | 5.8205 | 7.0616 | 14.1396 |
| Na | 0.5576 | 9.1339 | 17.5196 |
| Na | -4.9172 | 7.7179 | 20.8996 |
| **Na** | **-8.5147** | **3.3530** | **24.2796** |
| Na | -10.8572 | -1.8907 | 26.5159 |
| Na | 10.8572 | -1.8907 | 24.1841 |
| Na | 5.8205 | -7.0616 | 19.6604 |
| Na | 0.5576 | -9.1339 | 16.2804 |
| Na | -4.9172 | -7.7179 | 12.9004 |
| Na | -8.5147 | -3.3530 | 9.5204 |
| Na | -8.8594 | 2.2923 | 6.1404 |
| Na | -5.8205 | 7.0616 | 2.7604 |
| Na | -0.5305 | 9.5858 | -0.4433 |
